# Inference of VEGFR2 dimerization kinetics on the cell surface by integrating single-molecule imaging and mathematical modeling

**DOI:** 10.1101/2025.06.03.657760

**Authors:** Jaime Guerrero, Zachariah Malik, FNU Bilal, Soma Jana, Aparajita Dasgupta, Khuloud Jaqaman

## Abstract

Inter-receptor interactions play a key role in receptor signaling, which is the first step in cell signaling in response to external stimuli. In the case of Vascular Endothelial Growth Factor Receptor 2 (VEGFR2), dimerization is necessary for activation. VEGFR2 undergoes reversible interactions also in the absence of ligand. For a quantitative understanding of transmembrane signal transduction, it is necessary to quantify the interaction kinetics of VEGFR2 on the cell surface. Live-cell single-molecule imaging (SMI) has the powerful ability to capture receptor interaction events in their native cellular environment with high spatiotemporal resolution. However, it reveals these interactions for only the labeled subset, which is a small fraction of the full population of receptors. We have previously shown that mathematical modeling, combined with SMI data, offers a route to compensate for this lost information and infer the population-level receptor interaction kinetics from SMI data. Here, we applied this approach to VEGFR2, both wildtype full length VEGFR2 (FLR2) and a truncated mutant consisting of its extracellular and transmembrane domains (ECTM), which has been shown to exhibit reduced homotypic interactions in the absence of ligand. We developed a stochastic mathematical model mimicking VEGFR2 diffusion and interactions and determined the unknown model parameters through a combination of direct experimental measurements and stochastic model calibration. We found that a model of dimerization was sufficient to describe VEGFR2 interactions in the absence of ligand. While ECTM was primarily monomeric, FLR2 exhibited a substantial fraction of dimers. Our inference revealed that the difference between FLR2 and ECTM was primarily in the dimer association rate constant, which was about an order of magnitude lower for ECTM than for FLR2. To our knowledge, this is the first time that the interaction kinetics of VEGFR2 have been calculated in live cells.

**Author summary:** Interactions between receptors on the cell surface play a large role in the initiation and modulation of cell signaling. For example, the receptor VEGFR2 in vascular endothelial cells must dimerize as a first step toward signaling. However, many interactions, including VEGFR2 dimerization, also take place in the absence of ligand stimulation. Therefore, elucidating when and where receptor interactions take place, and what their kinetics are, is critical for understanding receptor and cell signaling, especially on a quantitative level. Determining interaction kinetics is particularly challenging, given that membrane proteins do not express well in vitro, and are thus not amenable to traditional biophysical methods that quantify molecular interactions. In addition, the membrane and cellular environment is much more complex than an in vitro setting. To address these challenges, we combine single-molecule imaging, stochastic mathematical modeling, and rigorous model calibration to derive the dimerization kinetics of VEGFR2 (both full-length and a truncation mutant with reduced dimerization) on the surface of live cells. To our knowledge, this is the first time that the interaction kinetics of VEGFR2 have been calculated in live cells. Our approach is generalizable to many other cell surface proteins, opening an avenue for quantifying molecular interaction kinetics in situ.

## Introduction

Cell surface receptors exhibit a high degree of dynamic organization, including myriad homotypic and heterotypic interactions [1-6]. There is mounting evidence that these interactions play a critical role in regulating receptor function and signaling [1-6]. However, we know little about the kinetics of these interactions, fundamental information for a quantitative, mechanistic understanding of receptor and cell signaling. This lack of quantitative characterization is largely because transmembrane proteins are not amenable to purification for traditional biophysical studies in vitro. More problematically, the in vitro solution environment is very different from the lipid bilayer environment in which cell surface receptors reside. Studies have shown that receptor interactions in the plasma membrane can have different energetics and kinetics from the interactions observed in vitro [7-12].

We previously developed an integrated single-molecule (SM) imaging (SMI) and computational modeling approach, termed “Framework for the Inference of in Situ Interaction Kinetics” (FISIK), to derive molecular interaction kinetics in their native cellular environment [13]. Briefly, SMI – in its many flavors – has the power to capture individual receptor interaction events in their native environment [14-21]. However, to achieve SM sensitivity, often only a small fraction of receptors is observed, and thus only a small fraction of interaction events is captured. To compensate for this missing information in SMI experiments, FISIK combines SMI data with stochastic mathematical modeling to infer, from the observed subset of interactions, the interaction kinetics of the full population of receptors. The main tenet is that, by incorporating experimental data acquisition artifacts into the model-generated data (such as substoichiometric labeling), the artifacts would be factored out when matching model-generated and experimental data, allowing us to learn the interaction kinetics of the true, underlying receptor system.

Here, we extended and applied FISIK to the cell surface receptor Vascular Endothelial Growth Factor Receptor 2 (VEGFR2), a critical receptor on the surface of vascular endothelial cells (VECs), where its signaling is important for the formation and maintenance of blood vessels in normal physiology and disease [2, 22]. VEGFR2 dimerization is a necessary step for its activation and signaling [2, 23, 24]. VEGFR2 can also dimerize in the absence of ligand, leading to important interplay between the basal state dynamic organization of VEGFR2, its binding to ligand, and subsequent activation steps [16, 25, 26]. For our purposes, VEGFR2 dimerization in the basal state provided us with an ideal steady-state system to apply FISIK, without the concern for transients (as would be the case upon stimulation). In addition, while there is generally very little quantitative characterization of the interaction kinetics of cell surface receptors in their native environment, a previous FRET-based study estimated the dissociation constant (*K*_*d*_) of VEGFR2 dimerization in plasma membrane derived vesicles that we could compare to our results [25]. Moreover, a truncation mutant of VEGFR2, consisting of only its extracellular and transmembrane domains (ECTM), was shown to undergo less dimerization than wildtype full-length VEGFR2 (FLR2) in the absence of ligand [25, 27], providing us with a negative control for comparison to FLR2. In contrast to previous work, which yielded only *K*_*d*_ for FLR2 and ECTM [25], our current work reveals the differences in interaction kinetics, i.e., association and dissociation rate constants, underlying the difference in *K*_*d*_ between FLR2 and ECTM.

## Results and Discussion

### Single-molecule analysis yields the apparent interaction kinetics for the labeled subset of VEGFR2 on the surface of CHO cells

To capture homotypic interaction events of wildtype, full length VEGFR2 (FLR2) or its truncated mutant with reduced dimerization (ECTM) at the SM level, we (transiently) expressed Halo-FLR2 or ECTM-Halo in CHO cells (Fig. 1A). FLR2 fused to HaloTag at its C-terminus (like ECTM) could not express on the cell surface, thus we employed FLR2 fused to HaloTag at its N-terminus in our study. CHO cells lack endogenous VEGFR2 [25], thus limiting VEGFR2 expression (whether FLR2 or ECTM) to that introduced exogenously. To minimize overexpression artifacts, both constructs were under a truncated CMV promoter, yielding expression levels similar to endogenous VEGFR2 in VECs (1-10 molecules/μm^2^ [28]). Additionally, we verified that Halo-FLR2 was functional by measuring VEGFR2 phosphorylation upon treatment with VEGF (2 nM for 5 min) in CHO cells expressing Halo-FLR2, which we found to be comparable to endogenous VEGFR2 phosphorylation in VECs in response to the same VEGF treatment (Fig. S1).

**Figure 1.**
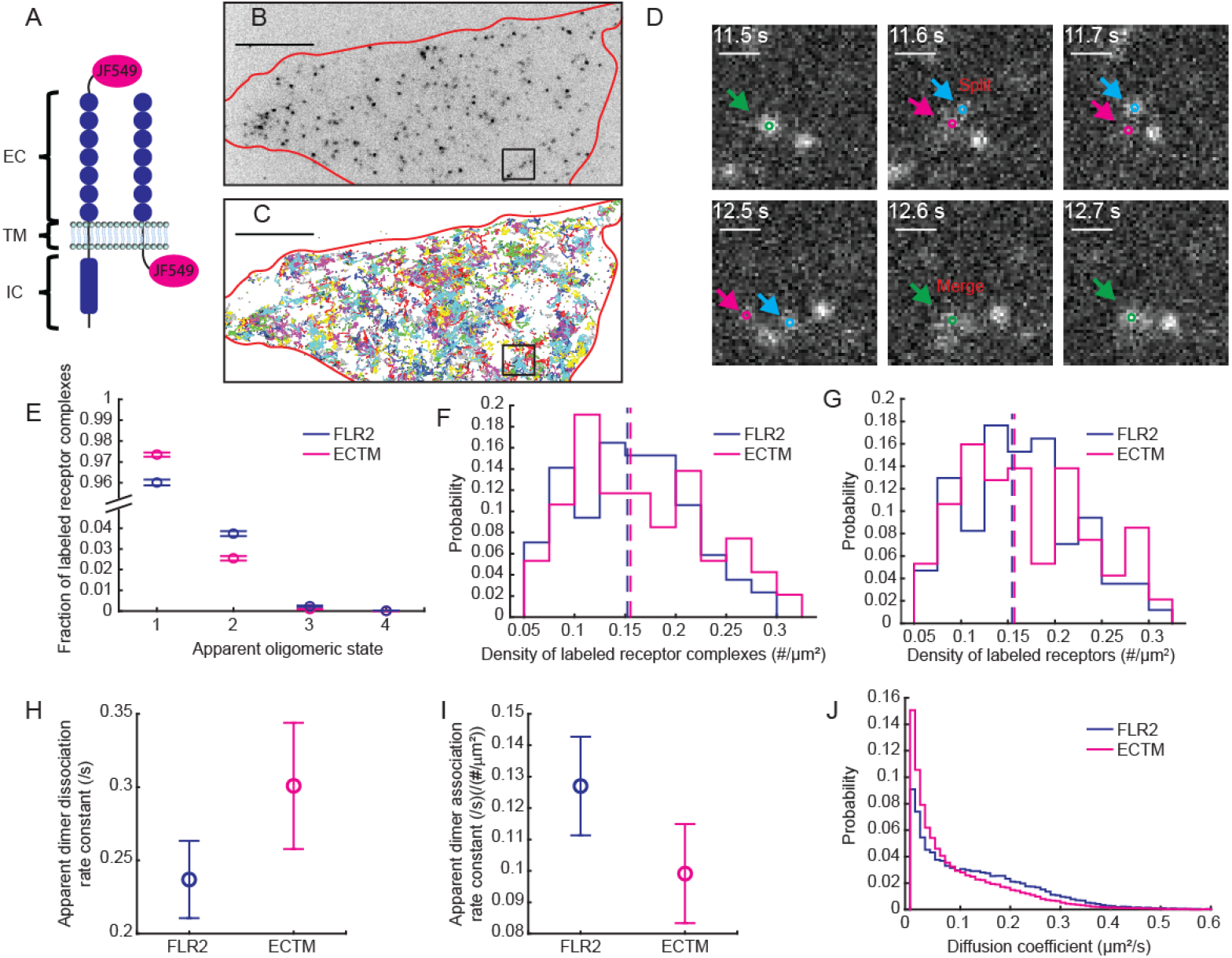
Live-cell single-molecule imaging yields the apparent homotypic interactions and their kinetics for the labeled subsets of FLR2 and ECTM on the cell surface. **(A)** Cartoon of FLR2 (full-length VEGFR2) and its ECTM truncation mutant, together with their HaloTag labeling. **(B)** Representative TIRFM image of Halo-FLR2 on the surface of CHO cells labeled at the single-molecule level with Halo-ligand conjugated to JF549, with cell outline in red. Halo JF549 labeling. Scale bar, 10 µm. The image is inverted for visual clarity. **(C)** Particle (i.e. labeled receptor complex) trajectories (with duration > 5 frames) from the cell shown in B, imaged for 30 s at 10 Hz. Trajectories are assigned random colors to help distinguish between them visually. Scale bar and red outline as in B. **(D)** Example of splitting and merging event from the boxed area in A-B. Scale bar, 10 µm. Orange arrow shows the particle when the two molecules are together, while cyan and magenta arrows show the two particles when separated. **(E)** Fraction of labeled receptor complexes, i.e., detected particles, in the indicated apparent oligomeric states, for FLR2 (blue) and ECTM (magenta). Circles show fractions from analyzing all imaged cells together. Error bars show the standard deviation of the fractions, estimated using Jackknife resampling. Note the cut in the y-axis. **(F)** Density of labeled receptor complexes per cell, shown as a distribution of individual cell measurements. Dashed vertical lines show mean values (0.157 for FLR2 and 0.156 for ECTM). **(G)** Density of labeled receptors per cell, shown as a distribution of individual cell values. Dashed vertical lines show mean values (0.164 for FLR2 and 0.171 for ECTM). **(H, I)** Apparent dimer dissociation **(H)** and association **(I)** rate constants. Circles and error bars as in E. **(J)** Diffusion coefficient distribution from all particles and all cells of each condition (FLR2 or ECTM). SMI dataset consisted of 85 (FLR2) and 94 (ECTM) cells, combined from 14 (FLR2) and 18 (ECTM) independent repeats.

After labeling Halo-FLR2 or ECTM-Halo with JF549-conjugated Halo ligand to achieve SM-level labeling, we imaged the transfected and labeled cells using total internal reflection fluorescence microscopy (TIRFM) to focus on receptors on the cell surface (Fig. 1B; Videos S1, S2). We acquired 10 Hz movies for 300 frames (30 s). We then detected and tracked the labeled receptors using u-track [29] (Fig. 1C; Videos S1, S2). U-track not only yielded receptor trajectories, but also their merging and splitting events (Fig. 1D; Videos S3, S4), which contained information about receptor interactions. Given the resolution limit, some of the captured merging and splitting events were most likely due to molecules passing by each other. As these events were expected to be short-lived, we refined the receptor trajectories by removing merge-to-split events lasting ≤ 2 frames (as well as those falling under several other criteria; see Materials and Methods). Note that some longer-lived events could also be due to resolution limit artifacts. This was compensated for in later analysis steps by incorporating experimental data acquisition artifacts into the model-generated data as part of the process of inferring the interaction kinetics of the full population from the labeled subset.

As a first step toward calculating the interaction kinetics of FLR2 or ECTM, we calculated the *apparent* interaction kinetics of the labeled subset of receptors. These kinetics are referred to as *apparent* because of the resolution limit, which creates “artifactual interactions”, and the subsampling inherent to SMI data, which renders many interactions invisible. Following the procedure described in [13], we first determined the apparent oligomeric state of each detected particle. Because we observed substantial variation in fluorophore intensity across our imaging area, leading to high overlap in the intensity distributions of different apparent oligomeric states (Note S1; Fig. S2), we took a conservative approach in the assignment of apparent oligomeric states. Specifically, we assumed that a particle was monomeric, unless merging and/or splitting events dictated otherwise [13]. Note that capturing merging and splitting events during tracking nevertheless followed strict local intensity constraints, where the intensity of the joint particle after a merge or before a split had to be roughly the sum of the intensities of the two merging or splitting particles, respectively [29]. For the remainder of this work, a detected particle (which could appear as monomeric, dimeric, etc.) will be referred to as a *receptor complex* (even if it is monomeric).

We found that the labeled subset of both FLR2 and ECTM appeared mostly as monomeric or dimeric, with very small fractions of FLR2 appearing as trimeric or tetrameric (Fig. 1E). Both imaged molecules had similar surface densities (Fig. 1F, G; see Suppl. Note S2, Eq. S3, for calculation of receptor density from receptor complex density), indicating that the differences between FLR2 and ECTM reflected differences in their underlying interaction kinetics. Although subtle, the difference in the apparent oligomeric state fractions between FLR2 and ECTM was consistent with ECTM undergoing less dimerization than FLR2. Of note, given the relatively small fraction of receptors that were labeled in our SMI experiments (10-15%; as calculated in later sections), this small difference in the oligomeric state fractions of the labeled subset reflected a large difference at the full population level (Note S2; Fig. S3).

From the merging and splitting events of each labeled receptor complex, we then calculated the apparent dissociation rate constants of the different apparent oligomeric state (Fig. 1H) [13]. Only apparent dimers had enough instances to allow this calculation. Combining the apparent dimer dissociation rate constant with the density of apparent monomers and dimers allowed us to then calculate the apparent dimer association rate constant (Fig. 1I) [13]. The differences between Halo-FLR2 and ECTM-Halo, namely ECTM having a lower apparent association rate constant and higher apparent dissociation rate constant, were again consistent with ECTM undergoing less dimerization than FLR2. The apparent oligomeric state densities and apparent interaction kinetics for the labeled subsets of FLR2 and ECTM constituted our intermediate (or summary) statistics describing the SMI data, to be used for comparing SMI data to simulated data to infer the full population homotypic interaction kinetics from the labeled subset.

### Inference of the interaction kinetics of the full population of VEGFR2 requires a well-constrained stochastic mathematical model and a scheme for generating synthetic SMI data

The core of the FISIK approach is to generate synthetic data that mimic SMI data – both the receptor system and the artifacts introduced via data acquisition – and then to compare the synthetic data to experimental SMI data while tuning the parameters of the mathematical model describing the receptor system until the two match (Fig. S4, Blocks IV and V) [13]. The main tenet is that, by incorporating experimental data acquisition artifacts into the synthetic data, the artifacts would be factored out when matching simulated and experimental data, allowing us to learn the true receptor system (mathematical model) parameters.

Generating synthetic data equivalent to SMI data was a multi-step process (Fig. S5A). First, we used particle-based reaction diffusion simulations to generate trajectories of the full population of diffusing and interacting receptors from a stochastic mathematical model (described next) [13, 30, 31]. Second, we introduced two layers of artifacts. The first layer was substoichiometric labeling, inherent to SMI experiments, leading to an underestimation of interactions [13]. The second layer involved making synthetic movies out of the simulated system, most importantly replacing each labeled receptor with a point spread function (PSF; approximated as a Gaussian with center = receptor position and standard deviation = 122 nm; see Materials and Methods), thus mimicking the resolution limit inherent to light microscopy, which overestimates interactions (Video S5). The synthetic movies also had fluorophore intensity, noise and motion blur properties similar to SMI movies (all derived directly from the SMI movies; see Materials and Methods). The synthetic movies were then detected, tracked and refined in the same manner as SMI movies (Fig. S5A; Video S5), thus introducing potential detection or tracking errors into the synthetic data, increasing their equivalence to the experimental SMI data.

The mathematical model mimicking VEGFR2 diffusion and interactions on the cell surface was an extension of that presented in [13]. In brief, receptors with density *ρ* diffuse in a 2D plane, where they are initially randomly placed. Each receptor (and, once interactions occur, receptor complex) has a diffusion coefficient *D* randomly sampled from the diffusion coefficient distributions observed experimentally for FLR2 or ECTM (Fig. 1J) (in [13], all receptors and receptor complexes in a simulation had the same diffusion coefficient). The receptors can associate with each other (one at a time) to form dimers, trimers, etc., up to a maximum allowable oligomeric state *N* (thus oligomeric state *n* = 1, 2, 3, …, *N*). Interaction kinetics are controlled by an association probability *p*_a_(*n*) and a dissociation rate constant *k*_off_(*n*) (which can be different for different *n*). To mimic substoichiometric labeling (layer 1 of artifacts), a fraction *p*_*l*_ of receptors is then labeled so that their trajectories and interactions are visible, while the rest remain invisible.

Among the above parameters, the maximum oligomeric state, association probability and dissociation rate constant were the interaction kinetics parameters of interest. To maximize the accuracy of their determination through model fitting to the SMI data [13], it was important to constrain the model by determining the remaining model parameters through orthogonal/independent experiments and analyses. For this, we pursued the following strategy: The diffusion coefficient distribution was estimated directly from the SMI data (Fig. 1J, Fig. S4, Block 1, Materials and Methods). The receptor density and labeled fraction were derived through complementary experiments and analyses (Fig. S4, Blocks II and III), described in the following two sections (Figs. 2, 3). Finally, for the maximum oligomeric state, we followed Occam’s razor and selected the model with the smallest maximum oligomeric state necessary to match the SMI data. All in all, this strategy highly constrained the model and increased the accuracy of determining the maximum oligomeric state, association probability and dissociation rate constant for VEGFR2 on the cell surface through model calibration with SMI data.

**Figure 2.**
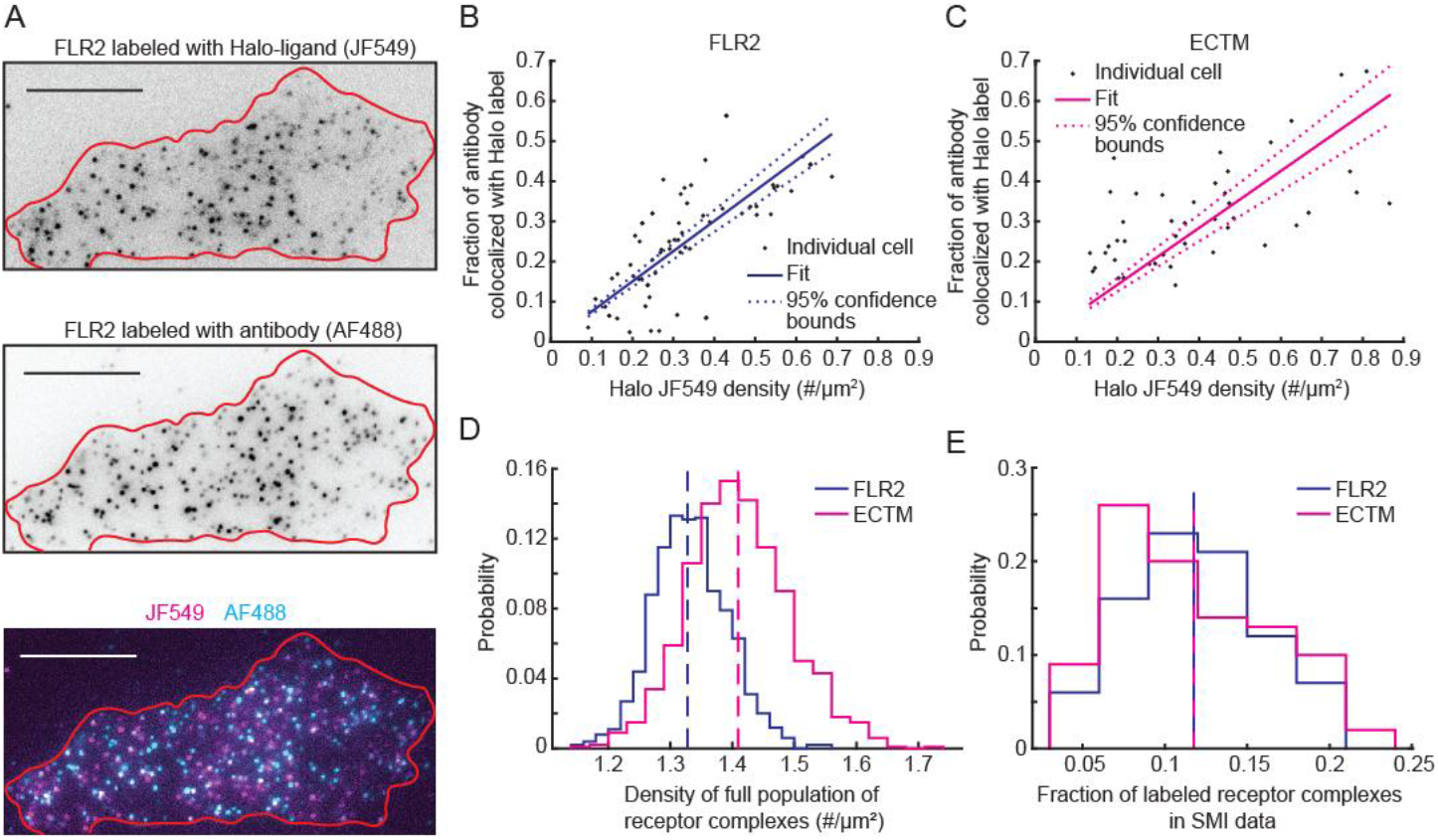
Two-color independent labeling of cell surface receptors yields the full population density and labeled fraction for FLR2 and ECTM. **(A)** Representative 2-color fixed-cell image of FLR2 in CHO cells, labeled with Halo-ligand and anti-VEGFR2 antibody. Cell outline is shown in red. The individual channel images are inverted for visual clarity. Scale bar, 10 µm. **(B, C)** Fraction of antibody detections colocalized with Halo label detections vs the density of halo label detections for FLR2 **(B)** and ECTM **(C)**. Each dot represents one cell. Solid and dashed lines show straight line fit (solid line) and its 95% confidence interval (dashed lines). **(D)** Density of the full population of receptor complexes, as derived from the inverse of the slope of the line fits in B and C. Distribution stems from randomly sampling the distribution of slopes, assuming a Gaussian distribution with mean and standard deviation estimated by the least squares fit. Dashed vertical lines show mean values (1.33 for FLR2 and 1.44 for ECTM). **(E)** Fraction of labeled receptor complexes per cell in the SMI dataset, shown as a distribution of individual cell measurements. Dashed vertical lines show mean values (0.118 for both FLR2 and ECTM). 2-color fixed cell dataset consisted of 65 (FLRs) and 44 (ECTM) cells, combined from 10 (FLR2) and 6 (ECTM) independent repeats. SMI dataset used for E same as that in Fig. 1.

**Figure 3.**
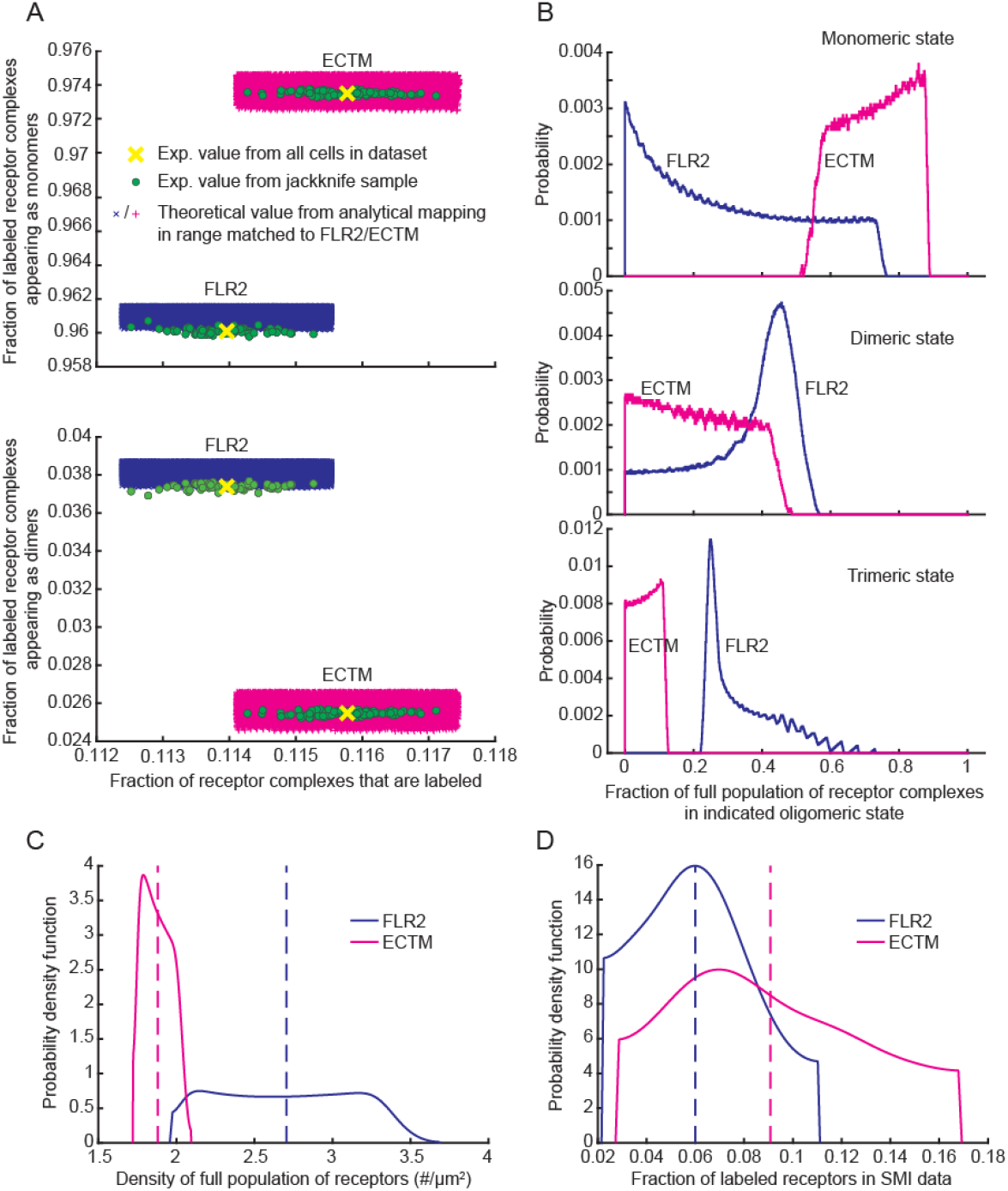
Analytical mapping yields the full population density and labeled fraction of FLR2 and ECTM on the cell surface from their receptor complex counterparts. **(A)** Range of theoretical values of fraction of labeled receptor complexes and fraction of apparent monomers and dimers among labeled receptor complexes matching the experimental measurements for FLR2 and ECTM. The theoretical values were obtained via analytical mapping between the full population and labeled subset of receptors and receptor complexes for a system that contains monomers, dimers and trimers (See Eqs. 3 and 4 in main text and Suppl. Note S2). Matching is shown as fraction of apparent monomers vs. fraction of labeled receptors (top) and fraction of apparent dimers vs. fraction of labeled receptors (bottom) for visual clarity. Fraction of apparent trimers is not shown as it is given by 1 - (fraction of apparent monomers + fraction of apparent dimers). **(B)** Fraction of monomers, dimers and trimers in the full population of receptor complexes underlying the matchings in A, for FLR2 and ECTM. **(C)** Density of the full population of receptors, as derived from combining the fractions of the different oligomeric states among the full population of complexes shown in B and the average density of the full population of receptor complexes shown in Fig. 2D. Distribution stems from the distribution of the fractions of the different oligomeric states shown in B. Dashed vertical lines show mean values (2.70 for FLR2 and 1.88 for ECTM). **(D)** Fraction of labeled receptors per cell in the SMI dataset, shown as a distribution of individual cell measurements. Dashed vertical lines show mean values (0.060 for FLR2 and 0.091 for ECTM). In C and D, the histograms were smoothened with a normal kernel (using the Matlab function “ksdensity”) to remove artificial jaggedness in the raw histograms resulting from the discrete sampling of the solution space. The smoothened histograms preserved the range of the raw histograms.

### Independent dual labeling analysis yields the surface density and labeled fraction of receptor complexes

To reduce the number of unknown model parameters and reduce the model degrees of freedom when fitting the model to SMI data, we sought to determine VEGFR2 (FLR2 or ECTM) receptor density and labeled fraction through complementary experiments and analyses. They were then fixed in the model, leaving only the interaction kinetics parameters of interest as unknown. This was a two-step process. First, using independent dual labeling analysis, we determined the density of *receptor complexes* on the cell surface and the fraction of *receptor complexes* that were labeled in the SMI experiments (this section). Then, using analytical mapping between the full population and labeled subset of receptors and receptor complexes (Suppl. Note S2), we mapped from density and labeled fraction of *receptor complexes* to those of *receptors* (next section).

To determine the density and labeled fraction of receptor complexes on the cell surface, we labeled Halo-FLR2 or ECTM-Halo with JF549-conjugated Halo ligand as in the SMI experiments and then, after cell fixation, we labeled them also with antibody targeting the extracellular domain of VEGFR2 (FLR2 or ECTM) (Fig. 2A). We then imaged the fixed cells using TIRFM, detected the imaged particles (i.e. labeled receptor complexes) in each channel, and, after correcting for registration shift between the two channels, calculated the fraction of antibody-labeled receptor complexes colocalized with JF549-labeled receptor complexes (using a colocalization radius of 3 pixels = 243 nm) [32, 33]. As these two labeling strategies were independent, this colocalization fraction was equivalent to the fraction of receptor complexes labeled with JF549 in the fixed cell setting, 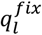. This fraction could be written as:

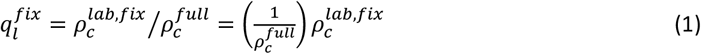

where 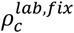 was the density of receptor complexes labeled with JF549 in the fixed cell setting, and 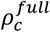 was the density of the full population of receptor complexes (labeled and unlabeled). Therefore, plotting the individual cell values of 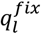 vs. 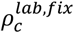 yielded 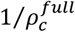, allowing us to estimate the density of the full population of Halo-FLR2 complexes and ECTM-Halo complexes (Fig. 2B-D).

As for the labeled fraction of receptor complexes *q*_*l*_, to estimate it directly for the live-cell SMI data, we divided the density of labeled receptor complexes per cell from the SMI data (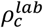; Fig. 1F) by the average density of the full population of receptor complexes estimated from the line fits (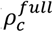; Fig. 2D):

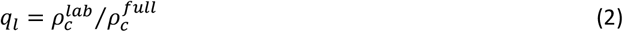

This yielded the distribution of the labeled fraction of receptor complexes per cell for Halo-FLR2 and ECTM-Halo in the SMI data (Fig. 2E).

### Full population-to-labeled subset analytical mapping yields the surface density and labeled fraction of receptors from their receptor complex counterparts

The above analysis yielded the full population density and the labeled fraction of *receptor complexes* on the cell surface (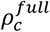 and *q*_*l*_, respectively). To determine the full population density and the labeled fraction of *receptors* on the cell surface (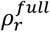and *p*_*l*_, respectively), as needed for our mathematical modeling, we employed analytical mapping between the full population and labeled subset of receptors and receptor complexes (Suppl. Note S2; Eq. 3 and 4 below). As derived in Suppl. Note S2, the fraction of labeled receptor complexes *q*_*l*_ (Fig. 2E) can be written as a function of the fraction of labeled receptors *p*_*l*_ and the fractions of the different oligomeric states in the full population of receptor complexes 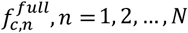 (Eq. S9 in Suppl. Note S2):

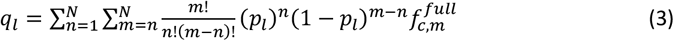

Therefore, determining the fraction of labeled receptors from the fraction of labeled complexes required knowledge of the fractions of the different oligomeric states in the full population of receptor complexes. However, our analysis of the SMI data thus far has only yielded the fractions of apparent oligomeric states among the labeled subset of complexes (Fig. 1E). Luckily, the same analytical mapping as above also provided equations that expressed the fractions of apparent oligomeric states among the labeled subset of complexes 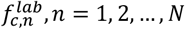 (Fig. 1E) as a function of their full population counterparts 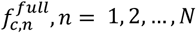, as well as the fraction of labeled receptors *p*_*l*_ (Eq. S8 in Suppl. Note S2):

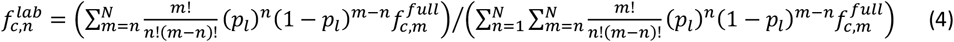

Together, Eqs. 3 and 4 provided us with *N* + 1 equations to determine *N* + 1 unknowns, namely the fraction of labeled receptors *p*_*l*_ and the fractions of the different oligomeric states in the full population of complexes 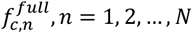. Combining 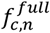 with the density of the full population of receptor complexes 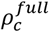 (Fig. 2D) through the equation (Eq. S3 in Suppl. Note S2):

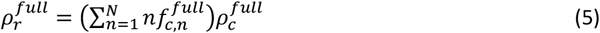

allowed us then to calculate the density of the full population of receptors 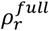.

Inverting Eqs. 3 and 4 to express the unknowns *p*_*l*_ and 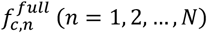 as functions of the knowns *q*_*l*_ and 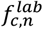 was not feasible. Instead, first we employed forward modeling to map combinations of *p*_*l*_ and 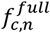 into *q*_*l*_ and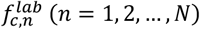, and then we matched the mapped *q*_*l*_ and 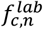 to their experimentally measured counterparts (Fig. 2E and Fig. 1E, respectively) in order to infer the underlying *p*_*l*_ and 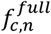. For the forward modeling, we scanned over all values of *p*_*l*_ and 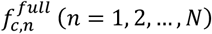 from 0 to 1 in steps of 0.001, such that 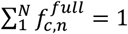.

As with the stochastic model of VEGFR2 diffusion and interactions, a critical step for the matching here was to determine the maximum oligomeric state of the system, *N*. Here too, we employed Occam’s razor and selected the smallest *N* necessary for matching. While *N* = 2 was sufficient to match the ECTM data, *N* = 2 could not match the FLR2 data. *N* = 3 was however able to match both (Fig. 3A), and thus we selected the *N* = 3 mapping for this analysis, for both FLR2 and ECTM. Note that this did not imply that the receptors formed trimers; at this stage, we did not yet factor out the effect of the resolution limit on the SMI data. Nevertheless, to estimate the full population density of receptors and their labeled fraction, it was necessary to use an *N* = 3 mapping.

From the *N* = 3 mapping and matching, we retrieved the fractions of the different oligomeric states in the full population of receptor complexes 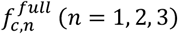 for FLR2 and ECTM (Fig. 3B). The solution spaces were a distribution of values for each fraction, because there were multiple combinations of *p*_*l*_ and 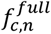 that led to similar *q*_*l*_ and 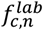. Nevertheless, there was a clear separation between FLR2 and ECTM, with ECTM having a higher fraction of complexes in the monomeric state while FLR2 had higher fractions of complexes in the dimeric and trimeric states. Importantly, combining these oligomeric state fractions for the full population of complexes with the average density of the full population of complexes determined from the line fit above (Fig. 2D) using Eq. 5 yielded the density of the full population of receptors 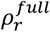 for FLR2 and ECTM (Fig. 3C). The density distributions stemmed from the distributions of oligomeric state fractions.

As for the fraction of receptors (FLR2 or ECTM) labeled in the SMI data, instead of taking its distribution directly from the solution space above, we calculated it per-cell. Specifically, we divided the density of the labeled subset of receptors per cell (Fig. 1G) by the average density of the full population of receptors (Fig. 3C), thus obtaining the labeled fraction of receptors per cell for each of FLR2 and ECTM (Fig. 3D). Not surprisingly, the labeled fraction of receptors (∼0.06 and ∼0.09 for FLR2 and ECTM, respectively) was smaller than the labeled fraction of receptor complexes (∼0.12) (Fig. S3B illustrates this trend using a monomer-dimer system as an example). Moreover, the difference between the labeled fraction of receptors and the labeled fraction of complexes was larger for FLR2 than for ECTM, reflecting the overall higher tendency of FLR2 to form complexes, while ECTM was overall more in the monomeric state.

The difference in the labeled fraction of receptors between FLR2 and ECTM (∼0.06 vs. ∼0.09) was also overall consistent with the concentrations of JF549 used for labeling them (6 nM for FLR2 vs. 12 nM for ECTM). Our guide at the time in choosing the concentration of JF549 was the density of labeled receptor complexes (i.e. detected particles) per cell, which could be calculated directly from the images without any analysis other than particle detection and cell mask delineation. Our choice of 6 nM for FLR2 and 12 nM for ECTM led to very similar labeled receptor complex densities (Fig. 1F). We originally attributed the need for different JF549 concentrations between the two molecules to the extracellular and intracellular location of the HaloTag in, respectively, Halo-FLR2 and ECTM-Halo. This might still be part of the reason, but this analysis shows that there were other reasons as well, namely the higher density of FLR2 on the cell surface (Fig. 3C) and the higher tendency of FLR2 to form complexes (Fig. 3B).

Regardless of the differences between FLR2 and ECTM, this analysis yielded distributions for the density of the full population of receptors and the fraction of receptors labeled in SMI experiments for the two molecules, which could be used for the mathematical model of FISIK.

### Dimerization model is sufficient to yield matching between model-generated SMI data and experimental VEGFR2 SMI data

With the above, we secured all parameters needed to model VEGFR2 diffusion and interactions on the cell surface, except for the unknown interaction parameters of interest, namely the maximum oligomeric state, the association probability per oligomeric state, and the dissociation rate constant per oligomeric state. These interaction parameters had to be determined by fitting the model to our SMI data via indirect inference [13, 34-36]. The maximum oligomeric state dictated the number of association probability and dissociation rate constant parameters to be determined (one each per oligomeric state > 1). Thus, we set the maximum oligomeric state in the model, and then determined the relevant parameters. Our approach here was to again follow Occam’s razor, i.e., start with the simplest model and increase the maximum oligomeric state one at a time, and then choose the model with the smallest maximum oligomeric state necessary for matching.

We started with a dimerization model. For a dimerization model, there were only two unknown model parameters, the dimer association probability and the dimer dissociation rate constant. The simpler model of no interactions was a special case of the dimerization model, with association probability set to zero. We generated simulated data (and then synthetic images and movies from the simulated data; see Materials and Methods, Fig. S5 and Video S5) using various combinations of dimer association probability (between 0.001 and 0.1) and dimer dissociation rate constant (between 0.01/s and 0.2/s), in addition to simulated data of no interactions (association probability = 0). Thus, we had a total of 51 parameter combinations, for each of which we generated 200 synthetic movies of duration 300 frames at 10 Hz, similar to our SMI data. We had one set of such simulations for FLR2 and one set for ECTM. The two molecules required separate simulations because of their different diffusion coefficient distributions, receptor densities and labeled fractions.

We then detected and tracked the synthetic movies and analyzed their apparent oligomeric states and interaction kinetics just like SMI data (Figs. S6-S9). Of note, even though these simulations contained only true monomers and dimers, they produced particles that appeared as trimers or tetramers (Figs. S7, S9), as observed in the experimental data. This is because the resolution limit of light microscopy was mimicked in our synthetic image generation.

To compare the synthetic data to experimental data quantitatively, we used the Mahalanobis distance between their intermediate statistics ([13]; Materials and Methods, Eq. 6). The intermediate statistics consisted of the densities of apparent monomers, dimers, trimers and tetramers in the system, in addition to the apparent dimer dissociation rate constant. The apparent dimer association rate constant was not used in the Mahalanobis distance, as it was redundant with the other statistics [13]. As with the experimental data (Fig. 1), the number of apparent trimers and tetramers in the simulated data was too small to calculate any apparent association or dissociation rate constants for those oligomeric states, and so we did not use them in the comparison. The apparent trimers and tetramers did however factor into the Mahalanobis distance in terms of their density.

The Mahalanobis distance yielded lowest-distance matches for both ECTM and FLR2 (Fig. 4), with simulation statistics similar to those measured experimentally (Figs. S6-S9). This implied that a dimerization model was sufficient to describe the homotypic interactions of FLR2 or ECTM on the cell surface, at least at the expression levels in our system, which was consistent with previous studies [27, 37]. Note that a dimerization model was also necessary, as the simulated data of no interactions did not match the SMI data for either FLR2 or ECTM. Interestingly, FLR2 was further away from the no interactions model than ECTM, again consistent with FLR2 undergoing more interactions than ECTM.

**Figure 4.**
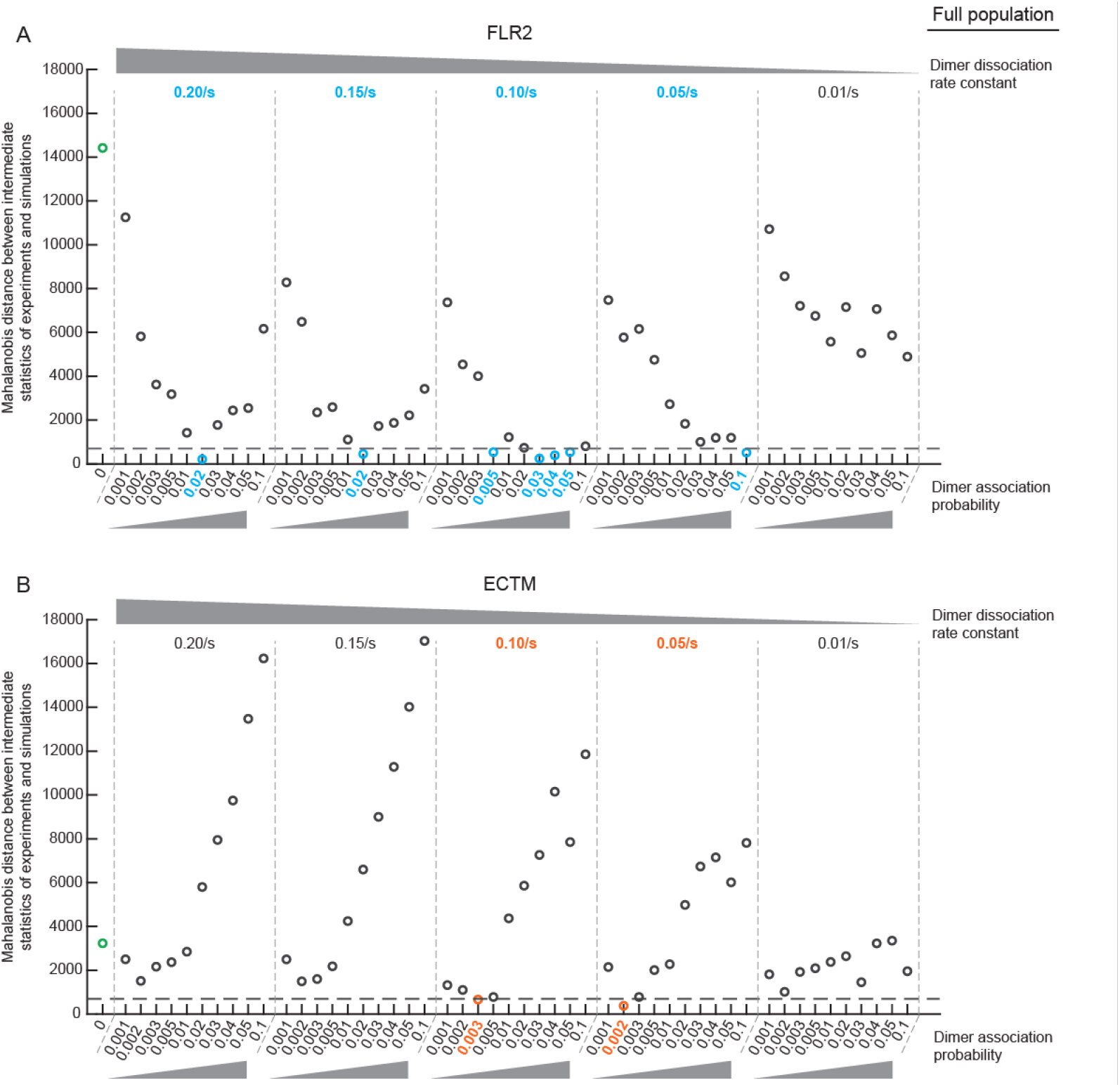
Comparison of SMI data to corresponding simulations identifies dimerization kinetics parameters that lead to matching between experiments and simulations. **(A, B)** Mahalanobis distance as a measure of dissimilarity between the intermediate statistics of simulated data and SMI data for FLR2 **(A)** and ECTM **(B)**. The intermediate statistics used for comparison are the densities of apparent monomers, dimers, trimers and tetramers among the labeled subset of receptors and the apparent dimer dissociation rate constant. The simulations, which were limited to dimerization, varied the dimer association probability and dissociation rate constant, with dimer association probability = 0 representing the special case of non-interacting molecules (shown as green circle). For visual clarity, the results are grouped by dissociation rate constant value (groups delineated by vertical dashed lines), within each of which the association probability is varied as indicated. Simulations matching the experimental data (Mahalanobis distance < 700; dashed horizontal line) for FLR2 or ECTM are highlighted by coloring their circles and parameter values in cyan in A or orange in B. Experimental intermediate statistics values are from Fig. 1. Simulated intermediate statistics are calculated from 200 simulations per interaction parameters combination (shown in detail in Figs. S6-S9).

### Indirect inference of full population interaction kinetics reveals that reduced dimerization of ECTM is primarily due to its lower association rate constant compared to FLR2

The Mahalanobis distance yielded lowest-distance matches for FLR2 and ECTM at very different model parameters (Fig. 4). For FLR2, the matches tended to be at higher association probabilities, mostly ≥ 0.02, while for ECTM the matching association probabilities were 0.002-0.003, i.e., an order of magnitude smaller than FLR2. The dissociation rate constant was not too different between the two molecules. These results suggested that the difference between ECTM and FLR2 was more in the tendency of the molecules to dimerize, and less in the stability of the dimer once it was formed. This could be because, without the intracellular domain, the extracellular and transmembrane domains do not possess the right conformations and/or orientations for dimerization.

To investigate this further and represent the association of VEGFR2 interactions in terms of an association rate constant, instead of association probability (which is specific to our model formulation [13]), we used the calibrated models to generate simulated data now at 100% labeling, i.e. the full population, and calculated the interaction kinetics of the full system directly from the simulations (without generating images and introducing artifacts). The calibrated model for each molecule (FLR2 or ECTM) was taken as the set of best matching models, specifically those with Mahalanobis distance < 700. 700 was taken as the threshold to yield the closest possible matches for each molecule without including too many peripheral matches. This yielded 7 matches for FLR2 and 2 matches for ECTM. These simulations were at 100 Hz and ran for 120 s to reduce discretization artifacts and capture the full spectrum of interaction durations (short-lived and long-lived).

These simulations identified significant differences between FLR2 and ECTM, consistent with ECTM undergoing much less dimerization than FLR2 (Fig. 5A-D). First, a much higher fraction of FLR2 than ECTM was dimeric (Fig. 5A; 12-60% for FLR2 vs. < 5% for ECTM). The fraction of dimeric FLR2 was consistent with previous studies [25]. Second, as suspected, the dimer association rate constant was the major difference between ECTM and FLR2 (Fig. 5C). Using these simulations and the association and dissociation rate constants that they yielded (Fig. 5B, C), we also calculated the dissociation constant of FLR2 and ECTM. Consistent with ECTM undergoing much less dimerization than FLR2, its dissociation constant was on average 14 mol/μm^2^, while the FLR2 dissociation constant was 2.6 mol/μm^2^ (Fig. 5D). The values of the dissociation constants estimated through our work were quite smaller than those estimated previously (35 mol/μm^2^ for FLR2 and 3200 mol/μm^2^ for ECTM [25]). These differences could stem from system differences, as the former measurements were made in plasma membrane-derived vesicles, which lack many regulatory elements, while the measurements in this work were made in intact membranes of live cells. Nevertheless, the dissociation constant of ECTM was about one order of magnitude larger than that of FLR2, consistent with ECTM exhibiting reduced dimerization.

**Figure 5.**
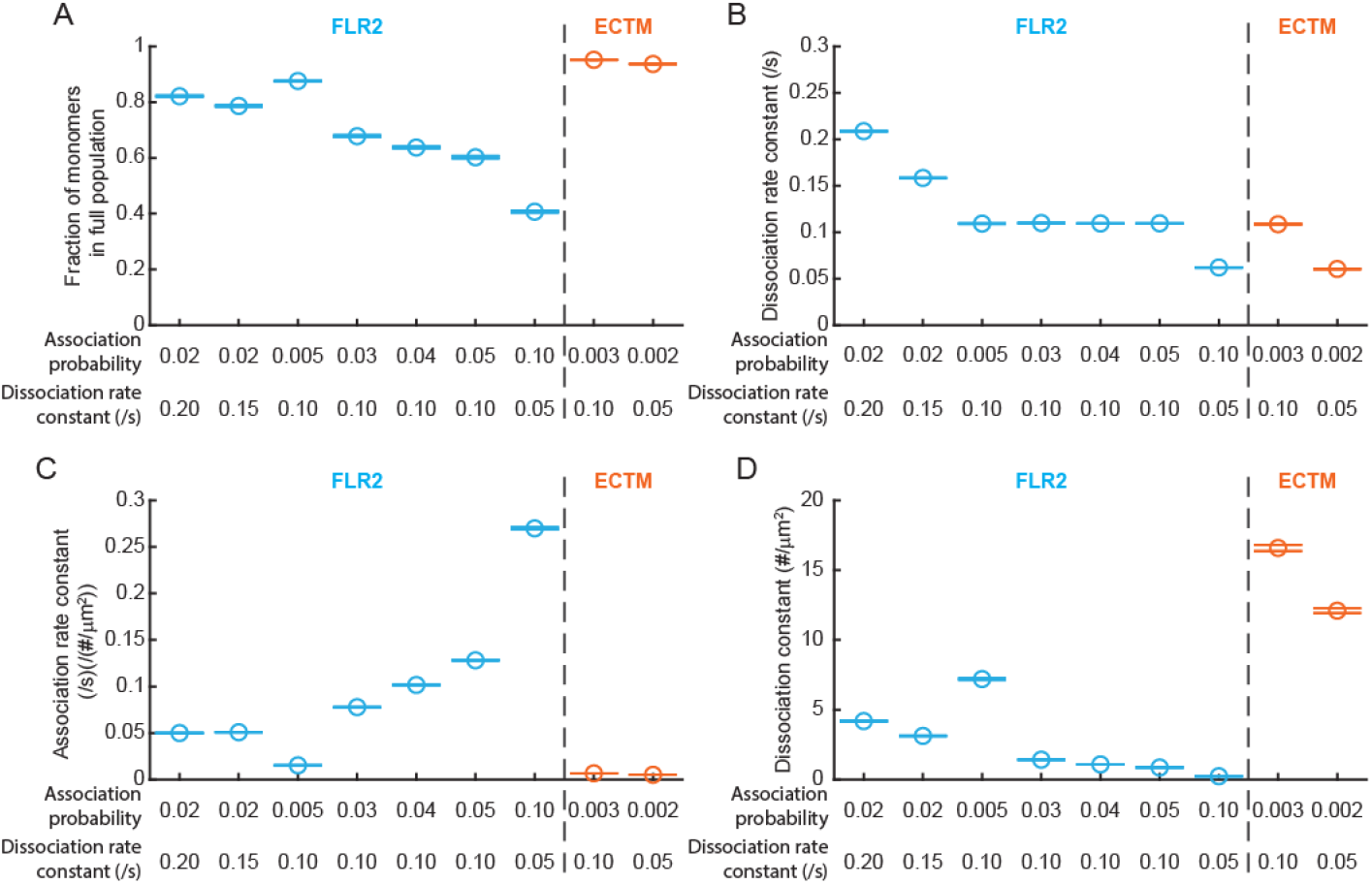
The identified interaction parameters predict the differences in dimerization kinetics underlying the lower dimerization tendency of ECTM compared to FLR2. **(A-D)** Predicted fraction of monomers among the full population of receptor complexes **(A)**, dimer dissociation rate constant **(B)**, dimer association rate constant **(C)** and dimer dissociation constant **(D)** for FLR2 (left in each panel; cyan) and ECTM (right in each panel, orange), as derived directly from simulations of FLR2 and ECTM with all receptors labeled (and no SMI artifacts). Interaction parameters used in the simulations correspond to the best-matching simulations in Fig. 4, and are indicated under each panel. Circles and error bars are the mean and standard deviation from 50 simulations (standard deviation obtained via Jackknife resampling).

In conclusion, we have determined in this work, for the first time to the best of our knowledge, the kinetics of VEGFR2 dimerization in an intact cell membrane. This information is critical for achieving a quantitative understanding of VEGFR2 signaling and regulation. The approaches developed in this work, whether the central approach of SMI combined with stochastic mathematical modeling, or the complementary approaches of independent dual labeling and analysis and the analytical mapping between the full populations and labeled subsets of receptors and receptor complexes, are general approaches that are applicable to any receptor system. They are also applicable beyond receptors, as long as the molecule(s) of interest can be labeled and imaged at the SM level. This approach will open avenues for deepening our quantitative understanding and characterization of receptor and other molecular interactions in the cell, which are the basis for cell signaling and cellular functions.

## Materials and Methods

### Plasmids

Plasmid encoding wildtype full-length VEGFR2 fused to HaloTag at its N-terminus (Halo-FLR2) was generated by VectorBuilder Inc. services (Chicago, IL). HaloTag (Promega, Madison, WI) was fused to the N-terminus of VEGFR2 via a EPTTEDLYFQSDN(AIA) linker and cloned into a pRP[EXP] vector containing an IL6 secretion signal [38]. A truncated CMV promoter (CMV100), courtesy of the Danuser lab (UT Southwestern, Dallas, TX), was used to reduce protein expression levels.

Plasmid encoding a truncated VEGFR2 mutant, containing only the extracellular and transmembrane domains, fused to HaloTag at its C-terminus (ECTM-Halo) was assembled as follows: The ECTM-mCherry (pcDNA3.1) vector, courtesy of the Hristova lab (Johns Hopkins University, Baltimore, MD) [25], was linearized using MluI and PspOMI restriction enzymes. Two gBlocks (Integrated DNA Technologies, Coralville, IA), one containing the CMV100 promoter with flanking regions homologous to the linearized vector and the ECTM coding sequence, and another containing the HaloTag sequence with flanking regions homologous to the ECTM linker ((GGS)x5) and linearized vector backbone, were synthesized. The ECTM-linker sequence was amplified from the ECTM-mCherry vector using forward primer ATGGAGAGCAAGGTGCTGCTG and reverse primer CGGTGCACTTCCTCCACTTCC. NEBuilder HiFi DNA assembly (New England Biolabs, Ipswich, MA) was used to assemble the DNA fragments to the plasmid backbone using the recommended protocol.

### Cell culture and transient transfections

Chinese Hamster Ovary (CHO-K1, ATCC, CCL-61, Manassas, VA) cells were grown in F-12K culture medium (ATCC, 30-2004) supplemented with fetal bovine serum (FBS, Thermo Fisher Scientific, A56697-01, Waltham, MA) added to 10% final concentration and 0.1X of antibiotic-antimycotic (Thermo Fisher Scientific, MT30004CI). Cells were grown at 37°C + 5% CO2, and were passaged once reaching 70-90% confluence (every ∼48 h). Passaging was done by trypsinization (0.05% Trypsin-EDTA, Thermo Fisher, 25300-054) for ∼5 minutes followed by resuspension in complete F12K medium at a 1:20 – 1:10 dilution. For imaging experiments, 1.5×10^5^ cells in 2 ml of medium were plated on base/acid cleaned glass bottom dishes (14 mm glass diameter with glass thickness of 0.17 mm (#1.5), MatTek, P35G-1.5-14-C, Ashland, MA) pre-coated for 45 min with 10 µg/ml fibronectin (Millipore Sigma, F2006-2MG, Burlington, MA). After 24-30 h of plating, when cells were at 60-90% confluence, cells were transfected with Halo-FLR2 or ECTM-Halo using Lipofectamine 3000 (Thermo Fisher Scientific, 25300-054) following the manufacturer’s instructions for 6-well dishes using 1.5 µg of the plasmid of interest. Transfected cells were grown overnight at 37°C + 5% CO2, until labeling and imaging. In total, imaging experiments were performed ∼2 days after cell plating.

Human Telomerase-Immortalized Microvascular Endothelial cells (TIME cells, ATCC, CRL-4025) were used for VEGFR2 phosphorylation experiments. Cells were passaged in ATCC’s vascular cell basal medium (ATCC, PCS-100-030) supplemented with microvascular endothelial cell (MVEC) growth kit-VEGF (ATCC, PCS-100-041), 12.5 µg/mL blasticidine (Thermo Fisher Scientific, A11139-03) and 0.1X of antibiotic-antimycotic (Thermo Fisher Scientific, MT30004CI) for 48 h at 37°C + 5% CO_2_ until reaching 80-90% confluence. Two days before experiments, 3.75 × 10^4^ cells were seeded on fibronectin-coated dishes as described above.

### Halo labeling for live-cell single-molecule imaging

Plated CHO cells transfected with VEGFR2 constructs as described above (2 days post seeding, 1 day post transfection) were first washed once in wash buffer (HBSS (Thermo Fisher Scientific, 14025-092) + 1 mM HEPES (Thermo Fisher Scientific, 15630-080) and 0.1% NGS (Jackson Immuno Research, 005-000-121, West Grove, PA)) to remove culture medium, then incubated at 37°C for 15 min with fresh pre-warmed F12K medium with Janelia Fluor 549 (JF549) HaloTag ligand (Promega, GA1110). For the main SMI experiments (Fig. 1), JF549-Halo ligand concentration was 6 nM (FLR2) or 12 nM (ECTM). For the low labeling SMI experiments for intensity analysis (Fig. S2), JF549-Halo ligand concentration was 0.5 nM (FLR2) or 1 nM (ECTM). The medium was then replaced with ligand-free F12K medium and incubated for 15 min to allow cells to recover from the wash. Incubation medium was then removed and cells were washed in wash buffer once, followed by the addition of pre-warmed imaging buffer (wash buffer + 100mM Oxyflour (Oxyrase, OF-0005, Mansfield, OH), 200mM trolox (Millipore Sigma, 238813) and glucose (Millipore Sigma, G8769)) to reduce photobleaching during imaging. Cells were then taken immediately to image.

### Halo and antibody labeling for 2-color fixed-cell colocalization imaging

Plated CHO cells transfected with VEGFR2 constructs were labeled with JF549-Halo ligand as described in the previous section (6 nM for FLR2 and 12 nM for ECTM). After the final wash for Halo labeling, samples were fixed with a 4% Paraformaldehyde solution (Electron Microscopy Sciences, 15714, Hatfield, PA) made in HBSS and incubated for 15 min at RT. Samples were then washed three times for 5 min each at RT, followed by 30 min blocking in blocking buffer (1% BSA, 5% NGS in wash buffer). The samples were then incubated for 1 h with 1:1000 primary antibody against VEGFR2 (4B4; ThermoFisher, MA5-15556) in blocking buffer. After washing three times (5 min each), samples were incubated with 1:1000 secondary antibody conjugated to AlexaFlour 488 (goat anti-mouse; ThermoFisher, A11001) for 15 min at RT. Finally, after three subsequent washes (5 min each), dishes were either incubated with a pre-warmed imaging buffer (Oxyfluor 1%, Glucose 0.45%, Trolox 2 nM) to proceed with imaging, or kept in washing buffer overnight at 4°C before imaging the next day.

### Halo and antibody labeling for fixed cell phosphorylation imaging

Cells (2 days post seeding and, in the case of CHO cells, 1 day post transfection) were serum starved for 3 h. For serum starvation, CHO cells were incubated in serum-free F12K medium, while TIME cells were incubated in serum-free vascular cell basal medium. Cells were then stimulated with 2 nM VEGF-A_165_ (GenScript, Z02689, Piscataway, NJ) for 5 min, followed by a quick 1 ml wash with wash buffer. Cells were then fixed with a 4% Paraformaldehyde solution made in HBSS and incubated for 15 min at RT. Samples were then washed three times for 5 min each at RT.

The procedure for VEGFR2 labeling varied by cell type. In the case of CHO cells, Halo-FLR2 was labeled during the serum starvation period (before VEGF addition and fixation) as follows. After 2.5 h of serum starvation, transfected CHO cells were quickly washed with 1 ml wash buffer, then incubated with 12 nM JF549-Halo ligand in serum-free F12K for 15 min, immediately followed by a 15 min incubation with serum-free F12K medium without Halo ligand. This yielded a total serum starvation time of 3 h. In the case of TIME cells, endogenous VEGFR2 was labeled after cell fixation. Here, TIME cells were blocked for 30 min in blocking buffer (1% BSA, 5% NGS in wash buffer), and then were incubated for 1 h with 1:1000 primary antibody against VEGFR2 (4B4; ThermoFisher, MA5-15556) in blocking buffer. Cells were then washed three times for 5 min each. Then, they were incubated for 15 min at RT with 1:1000 secondary antibody conjugated to AlexaFluor 568 (goat anti-mouse; ThermoFisher, A11004). Cells were then washed again three times for 5 min each.

After labeling VEGFR2, both CHO and TIME cells were permeabilized with cold permeabilization buffer (0.1% Triton X-100 in HBSS) (Thermo Fisher Scientific, BP151-100) for 1 min, followed by washing three times for 5 min each. Cells were then blocked for 30 min in blocking buffer. Blocking buffer was then replaced with 1:100 anti-phospho-VEGFR2 antibody (19A10, Cell Signaling Technology, 2478, Danvers, MA) diluted in blocking buffer to detect phosphorylated VEGFR2. Cells were then washed three times for 5 min each, followed by incubation with 1:1000 secondary antibody conjugated to AlexaFluor 647 (Goat anti-rabbit; ThermoFisher, A32733) for 15 min at RT. Finally, after three subsequent washes (5 min each), dishes were incubated with either a pre-warmed imaging buffer (Oxyfluor 1%, Glucose 0.45%, Trolox 2 nM) to proceed with imaging, or kept in washing buffer overnight at 4°C before imaging the next day.

### Live-cell single-molecule imaging

Live CHO cells (transfected with VEGFR2 constructs and labeled as described above) in imaging buffer were imaged at 37°C using an Olympus IX83 TIRF microscope equipped with a Z-Drift Compensator and a UAPO 100X/1.49 NA oil-immersion TIRF objective (Olympus, Tokyo, Japan). The microscope was equipped with an iXon 888 1k × 1k EMCCD Camera (Andor; Oxford Instruments, Oxfordshire, England). With an additional 1.6X magnification in place, the pixel size in the recorded image was 81 nm × 81 nm. Using the Olympus cellSens software, excitation light of 561 nm from an Olympus CellTIRF-4Line laser system was directed to the sample by a TRF8001-OL3 Quad-band dichroic mirror. Laser power at sample position (in the widefield illumination configuration) was ∼5.3 mW for the 561 laser. Fluorescence was collected, filtered with emission filters of ET520/40m, ET605/52m, and ET705/72m (Chroma), and projected onto different sections of the camera chip by an OptoSplit III 3-channel image splitter (Cairn Research, Faversham, UK). The penetration depth was set to 90 nm via the cellSens software (Olympus). Camera EM gain was set to 100 for all acquisitions. Videos were acquired with MetaMorph (Molecular Devices, San Jose, CA) in the stream acquisition mode, using the 561 channel. These were acquired at 10 Hz for 30 s for the main SMI experiments (Fig. 1) or for 100-250 s for the low labeling SMI experiments for intensity analysis (Fig. S2). The 561 channel was triggered to remain open (i.e., continuous illumination) for the entire 300 frames. Temperature and humidity were maintained using an environment chamber (Okolab, Otaviano, Italy), maintaining cell viability for the duration of the experiments.

For every SMI movie, a brightfield (BF) and/or interference reflection microscopy (IRM) snapshot of the imaged cell region was also acquired, to aid with manual delineation of the region of interest (ROI) mask for the ensuing analysis.

### Fixed-cell imaging

For both the 2-color colocalization and phosphorylation experiments, fixed CHO cells (transfected with VEGFR2 constructs and treated and/or labeled as described above) in imaging buffer were imaged at 37°C using the Olympus IX83 TIRF microscope described in the previous section. For these experiments, the two channels were excited and recorded sequentially, starting with 561 followed by 491 (2-color colocalization) or 640 (phosphorylation), with an exposure time of 99 ms each. Laser powers at sample position (in the widefield illumination configuration) were ∼1.3, ∼5.3, ∼4.1 mW for 488, 561 and 640 respectively.

For every two-channel image, a brightfield (BF) and/or interference reflection microscopy (IRM) snapshot of the imaged cell region was also acquired, to aid with manual delineation of the region of interest (ROI) mask for the ensuing analysis.

### Bead preparation and imaging for registration shift correction in fixed cells

To acquire images of Tetraspeck beads (Thermo Fisher Scientific, T7279) for calculating the registration shift between the 2 channels in fixed cell images (561 and 491 or 561 and 640), Tetraspeck beads were suspended (1:100) in distilled water using a sonicator-water bath for 15 min. The mixed bead sample was then combined with 10 µl poly-L-Lysine (Newcomer supply, Middleton, WI) diluted 1 in 10 µl in nuclease free water, and then plated on a cleaned Mattek dish (as used for cells but without fibronectin coating) for 30 min at RT in the dark. The bead sample was imaged using the Olympus IX83 TIRF microscope described above. The penetration depth was set to 90 nm via the cellSens software (Olympus). Laser powers at sample position (in the widefield illumination configuration) were ∼0.62, ∼1.8 and ∼1.4 mW for 488, 561 and 640 respectively.

### Cell ROI mask

For both fixed and live-cell data, cell ROI masks were hand-drawn using the u-track package (https://github.com/DanuserLab/u-track). Cell outlines were hand drawn in either the BF or IRM image, with reference to the associated fixed-cell or SMI data.

### SMI data: Particle detection and tracking

Particle detection and tracking for both real SMI movies and synthetic images were performed using u-track [29] (https://github.com/DanuserLab/u-track). Particle detection used the “Gaussian mixture-model fitting” algorithm. The particle localization precision in our SMI movies, estimated through the Gaussian least squares fit, was 14.6 ± 6.4 nm. The detection and tracking parameters, listed in the tables below (for non-default values), were optimized based on visual inspection of the detection and tracking results. In addition to the explicitly user-defined parameters, the number of frames used to calculate the intensity before and after merges and splits inside the tracking code (line 807 in costMatRandomDirectedSwitchingMotionCloseGaps) was changed from 2 to 10, in order to increase the stringency of capturing merging and splitting events. In the case of experimental data, only tracks within the cell ROI mask were retained for further analysis.

#### U-track detection parameters (Gaussian Mixture-Model Fitting)

Shown are the parameters with non-default values, based on u-track Version 2.2.1.

| Parameter | Value |
| --- | --- |
| Local maxima detection:<br>α-value for comparison with local background | 0.1 |
| Gaussian mixture-model fitting at local maxima:<br>Do iterative Gaussian mixture-model fitting | Yes |
| Gaussian mixture-model fitting at local maxima:<br>α-value for residuals test | 0.05 |
| Gaussian mixture-model fitting at local maxima:<br>α-value for amplitude test | 0.1 |
| Gaussian mixture-model fitting at local maxima:<br>α-value for distance test | 0.1 |

#### U-track tracking parameters

Shown are the parameters with non-default values, based on u-track Version 2.2.1.

***U-track tracking parameters.*** Shown are the parameters with non-default values, based on u-track Version 2.2.1.
| Parameter | Value |
| --- | --- |
| Overall:<br>Do segment merging | Yes |
| Overall:<br>Do segment splitting | Yes |
| Overall:<br>Maximum gap to close | 5 |
| Frame-to-frame linking:<br>Brownian search radius upper bound | 10 pixels |
| Frame-to-frame linking:<br>Multiplication factor for Brownian search radius calculation | 4 |
| Frame-to-frame linking:<br>Number of frames for nearest neighbor distance calculation | 6 |
| Gap closing, merging and splitting – general:<br>Brownian search radius upper bound | 10 pixels |
| Gap closing, merging and splitting – general:<br>Number of frames for nearest neighbor distance calculations | 6 |
| Gap closing, merging and splitting – general:<br>Scaling power in fast expansion phase | 0.25 |
| Gap closing, merging and splitting – general:<br>Gap length to transition from fast to slow expansion | 5 |
| Gap closing, merging and splitting – general:<br>Use nearest neighbor distance to expand Brownian search radius | Yes |
| Gap closing, merging and splitting – merging & splitting:<br>For an end/start, the possibility of merging/splitting is allowed only if there is no possibility of gap closing | Yes |
| Gap closing, merging and splitting – merging & splitting:<br>Min allowed | 0.75 |
| Gap closing, merging and splitting – merging & splitting:<br>Max allowed | Inf |
| Gap closing, merging and splitting – birth & death:<br>Birth and death cost | Derived from gap closing and merging/splitting cost formulae |
| Kalman filter functions:<br>Search radius for first iteration | 4 frames |

The tracks as output by u-track (for both experimental and simulated data) were then refined to remove spurious merging and splitting events, as follows: If a merge was followed by a split within ≤ 2 frames, both merge and split were removed. If a split was followed by a merge of the same two particles within ≤ 2 frames, both split and merge were removed. If a particle split and then disappeared within ≤ 2 frames, the split was removed. If a particle appeared and then merged with another particle within ≤ 2 frames, the merge was removed.

##### SMI data: Diffusion analysis

To construct the distribution of diffusion coefficients for simulating VEGFR2 (FLR2 or ECTM) diffusion and interactions, first the diffusion coefficient of each track in the SMI (experimental) data was calculated from its average square frame-to-frame displacement over its lifetime, taking into account its localization precision, as previously described [16, 39]. Then, a distribution of diffusion coefficients was constructed from these individual track measurements, such that each track’s contribution to the distribution was weighted by the track’s lifetime. Weighted contributions compensated for lifetime variation between tracks, and minimized bias in the distribution resulting from faster molecules getting tracked less well than slower molecules. Practically, weighting was done by repeating the diffusion coefficient of each track by the number of frames in the track when constructing the histogram of diffusion coefficients.

### SMI data: Calculation of apparent oligomeric state densities and fractions and of apparent association and dissociation rate constants

To compare between experimental and simulated data (Fig. 4), the apparent oligomeric state densities and fractions and the apparent association and dissociation rate constants for the labeled subset were calculated for all cells of a particular condition together (experimental data; Fig. 1E, H, I)) or for all simulations of a particular combination of model parameters together (simulated data; Figs. S6-S9). The uncertainty in the calculated statistics (error bars in Fig. 1E, H, I and in Figs. S6-S9) was estimated using Jackknife resampling [40], i.e., by taking out one cell/simulation at a time and calculating the statistics from the remaining cells/simulations. The error bars are the standard deviation among the Jackknife samples.

These same procedures were also followed for the full population simulations in Fig. 5, although in that case the calculated statistics were for the full population and without SMI artifacts; thus they were not “apparent.” In the below description, we will drop using the term “apparent”, with the understanding that if the statistics are calculated from the labeled subset, then they are apparent.

The statistics were calculated as described in [13] (https://github.com/kjaqaman/FISIK). In brief, the oligomeric state of each detected particle was determined using constrained least squares minimization (Eqs. 2 and 3 in [13]). Practically, to impose the rule that a particle was monomeric unless merging and/or splitting events dictated otherwise (because of the large fluorophore intensity variation, as discussed in Results), the intensity of an individual fluorophore in Eq. 2 in [13] was set to a very large number. Having the oligomeric state of each particle at each time point, the density of receptor complexes (i.e., particles) of oligomeric state 1, 2, etc. per time point was calculated by dividing the number of particles of oligomeric state 1, 2, etc. by the cell mask area (both summed over all cells/simulations in sample). This was then averaged over all time points in the SMI movies/simulations. The total density of receptor complexes regardless of oligomeric state and the fraction of each oligomeric state were then calculated from the time-averaged oligomeric state densities using Eqs. S1 and S2 in Suppl. Note S2.

Treating transitions between oligomeric states as a Markov process, the dissociation rate constant per oligomeric state > 1 was calculated from the oligomeric state history of each detected particle and the associated merging and splitting events yielded using Eq. 4 in [13]. Finally, assuming steady state, the association rate constant per oligomeric state > 1 was calculated from the dissociation rate constant and oligomeric state densities using Eq. 5 in [13].

To calculate labeled receptor complex (i.e., particle) density on a per-cell basis (Fig. 1F), the same density calculation procedures were followed as above, but treating each cell as its own sample. The labeled receptor density on a per-cell basis (Fig. 1G) was then calculated from the complex density on a per-cell basis using Eq. S3 in Suppl. Note S2.

### SMI data: Comparison of intermediate statistics using Mahalanobis distance

To compare the intermediate statistics of experimental and simulated data, we used the Mahalanobis distance [13], defined as:

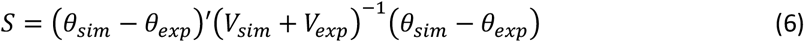

*θ*_*sim*_ and *θ*_*exp*_ were the vectors of intermediate statistics for simulated and experimental data, respectively. They consisted of the densities of apparent monomers, dimers, trimers and tetramers, as well as the apparent dimer dissociation rate constant, calculated from the full dataset (multiple cells or multiple simulations) for each condition, as described in the previous section. *V*_*sim*_ and *V*_*exp*_ were the variance-covariance matrices of the simulated and experimental intermediate statistics, respectively, estimated using the Jackknife, as described in the previous section.

### Fixed cell data: Particle detection and channel registration correction

Particle detection in fixed-cell and Tetraspeck bead images was performed using “point-source detection” in u-track [29] (https://github.com/DanuserLab/u-track). Default parameter values were used, except for the α-value used for selecting objects by comparing the fitted Gaussian amplitude to the local background noise distribution. The α-values for 640, 561 and 488 channels were, respectively, 0.005, 0.045 and 0.04 for the cell data and 0.001, 0.000001 and 0.001 for the bead data. These values were optimized via manual inspection, with the goal of minimizing both false-positives (superfluous detections) and false-negatives (missed particles).

To increase the accuracy of the ensuing colocalization analysis, the coordinates of detected particles in the 640 (phosphorylation experiment) or 488 (2-colocalization experiment) channel were aligned relative to the 561 channel. This was achieved by calculating a registration shift between the three channels, using Tetraspeck bead images. In brief, the Tetraspeck bead detections in the three channels were matched between channels using the Linear Assignment Problem [29], using a search radius of 9 pixels between 491 and 561 and a search radius of 5 pixels between 640 and 561. The 561 channel was taken as the reference. The matchings from multiple images acquired on the same day (3-5 images) were then utilized together to increase imaged area coverage. Using the matched bead detections, the registration shifts in x and y (Δx and Δy) as a function of position (x and y) were fit using a quadratic polynomial, deemed visually as the most appropriate fit. With this fit, the detections in the fixed-cell multi-channel images were then aligned prior to colocalization analysis.

### Fixed cell data: Two-color colocalization and related analyses

Colocalization analysis of the fixed-cell images (after channel registration) was performed as described in [32, 33]. This object-based colocalization analysis yielded the fraction of target objects colocalized with reference objects as a measure of colocalization. A target object was considered to be colocalized with reference objects if the distance between the target object and its nearest neighbor reference object was smaller than a user-defined threshold (“colocalization radius”, taken as 3 pixels = 243 nm).

In the case of the independent dual labeling analysis (Fig. 2), the antibody detections were the target and the Halo-label detections were the reference. After obtaining the colocalized fraction per cell, the plot of the fraction of antibody colocalized with Halo-label vs. the density of Halo-label per cell was fitted with a straight line passing through zero, in order to obtain the full population density of receptor complexes (Eq. 1). The fit was performed via least squares using the Matlab function “fitlm”, which also yielded the 95% confidence interval of the line slope (shown in Fig. 2B, C) and the slope’s standard deviation, the latter used for sampling to obtain a distribution for the full population density of receptor complexes (Fig. 2D).

In the case of the phosphorylation experiments (Fig. S1), VEGFR2 detections were the target and phospho-VEGFR2 detections were the reference. The colocalized fractions between stimulated and unstimulated cells were compared for each of Halo-FLR2 in CHO cells and endogenous VEGFR2 in TIME cells using a two-sample t-test.

### Simulation of VEGFR2 diffusion, interactions and sub-stoichiometric labeling

These simulations were based on those described in [13], with the added feature that receptors within a simulation had individual diffusion coefficients drawn from the distribution of FLR2 or ECTM diffusion coefficients measured experimentally (Fig. 1J) (in the original simulations, all receptors in a simulation had the same diffusion coefficient). Here we list details specific to the simulations in this work. Each parameter combination (association probability and dissociation rate constant) was represented by 200 simulations (for the model calibration simulations; Fig. 4) or 50 simulations (for the full system simulations after identifying the matching model parameters; Fig. 5). For each simulation, the receptor density and labeled fraction were randomly chosen from their experimentally-derived distributions (Fig. 3C, D). The simulation area was taken as 25 × 25 μm^2^ (with reflective boundaries), so that there were no boundary effects on the simulated system [13]. Receptors were randomly placed in the simulation area, initially all as monomers, and each simulation started with a 10 s initialization time for the system to reach steady state. Simulations were then run for 30 s (for the model calibration simulations, like the SMI data) or 120 s (for the full system simulations after identifying the matching model parameters). Simulation time step was 0.01 s.

### Generation of synthetic single-molecule movies from simulations

Synthetic single-molecule movies were generated from the above-described simulations by placing the molecules in a pixelated image, with pixel size = 81 nm like the experimental data. The synthetic SM movies were generated in the following manner to much as possible.

#### Point spread function (PSF), motion blue, and sampling rate

A Gaussian with standard deviation = 1.5 pixels (= 122 nm), approximating the PSF of a single fluorophore, was placed at the location of each molecule. Movies were first generated at 100 Hz (simulation time step of 0.01 s), and then every 10 images were averaged to make the final movies at a frame rate of 10 Hz, like the experimental SMI movies. This time-averaging introduced motion blur for the faster-moving molecules, mimicking the experimental images. With this, the effective Gaussian standard deviation of simulated particles, stemming from both the PSF and motion blur, was 1.58 pixels (= 128 nm), equal to that measured in the experimental SMI movies.

The effective Gaussian standard deviation in both the experimental and synthetic SMI data was determined by fitting a Gaussian with variable standard deviation to isolated particles in the experimental and synthetic SM images, respectively, using the Gaussian mixture-model fitting algorithm in u-track [29] (https://github.com/DanuserLab/u-track).

#### Fluorophore intensity fluctuations

To capture fluorophore intensity fluctuations between frames, the amplitude of the Gaussian per molecule was simulated to follow a normal distribution with a specified mean and standard deviation. The mean of the single-fluorophore intensity distribution was related to the signal-to-noise ratio, defined next. Given a particular mean, the standard deviation of the single-fluorophore intensity distribution was then defined as 0.25 x the mean, mimicking single-fluorophore intensity fluctuations in the experimental SMI data (Fig. S2E).

The single-fluorophore intensity standard deviation-to-mean ratio in the SMI data was calculated per single-fluorophore trace by taking its intensity standard deviation over its lifetime and dividing it by its mean intensity over its lifetime. Traces were deemed to represent individual fluorophores if they satisfied the following criteria: (i) trace disappeared before the end of the movie, (ii) trace had 0 photobleaching steps, (iii) trace had a mean intensity less than the mean photobleaching step size (Suppl. Note 1; Fig. S2B); and (iv) trace lifetime was at least 10 time points. This yielded the single-fluorophore intensity standard deviation-to-mean ratio in the experimental SMI data, which was on average 0.24 for FLR2 and 0.22 for ECTM (Fig. S2E).

#### Signal-to-noise ratio (SNR)

Underneath the placed molecules, the image background was simulated as white noise, following a normal distribution with a specified mean and standard deviation. The mean amplitude of the Gaussian per molecule (representing a single fluorophore), together with the image background mean and standard deviation, were chosen such that the signal-to-noise ratio (SNR) in the final movies (at 10 Hz) was ∼5.5, similar to that of the experimental SMI data.

The SNR of experimental data (per movie) was calculated as follows: First, the intensity of the detected particles at their center position was taken. Second, background intensity was taken as the intensity of all pixels in the ROI outside of a 4-pixel radius around each detected particle. The SNR for each movie was then calculated as (particle intensity - mean background intensity)/(standard deviation of background intensity). The average SNR for all the movies was 5.5 for ECTM and 6.8 for FLR2. The lower SNR value was chosen when creating simulations.

## Supporting information

Supporting material consists of two notes, 9 figures and 5 videos.

Note S1: Step photobleaching analysis.

Note S2: Analytical mapping between full populations and labeled subsets of receptors and receptor complexes.

Figure S1: Validation of Halo-FLR2 construct.

Figure S2: Analysis of fluorophore intensity properties in SMI data.

Figure S3: Analytical mapping between full population and labeled subset in a system that contains monomers and dimers.

Figure S4: Flowchart of experiments, analyses and data flow to determine the kinetics of VEGFR2 homotypic interactions on the cell surface.

Figure S5: Pipeline of generating synthetic single-molecule data of VEGFR2 diffusion and interactions to compare to experimental SMI data.

Figure S6: Statistics describing experimental and simulated FLR2 interactions, Part I.

Figure S7: Statistics describing experimental and simulated FLR2 interactions, Part II.

Figure S8: Statistics describing experimental and simulated ECTM interactions, Part I.

Figure S9: Statistics describing experimental and simulated ECTM interactions, Part II.

Video S1. Live-cell single molecule imaging and particle tracking of Halo-FLR2 in CHO cells.

Video S2. Live-cell single molecule imaging and particle tracking of ECTM-Halo in CHO cells.

Video S3. Close-up view of example particle merging and splitting events for Halo-FLR2.

Video S4. Close-up view of example particle merging and splitting events for ECTM-Halo.

Video S5. Synthetic single-molecule movie.

## Video legends

**Video S1. Live-cell single molecule imaging and particle tracking of Halo-FLR2 in CHO cells**. Representative 10 Hz/30 s TIRFM movie of Halo-FLR2 on the surface of CHO cells labeled at the single-molecule level with Halo-ligand conjugated to JF549. Same cell shown in Fig. 1B, C. Shown are the raw movie (top) and the particle tracks overlaid on the movie (bottom). Tracks are color-coded randomly with 7 different colors for ease of visualization. Image size is 27.6 × 64.8 µm^2^.

**Video S2. Live-cell single molecule imaging and particle tracking of ECTM-Halo in CHO cells**. Representative 10 Hz/30 s TIRFM movie of ECTM-Halo on the surface of CHO cells labeled at the single-molecule level with Halo-ligand conjugated to JF549. Shown are the raw movie (top) and the particle tracks overlaid on the movie (bottom). Tracks are color-coded randomly with 7 different colors for ease of visualization. Image size is 27.6 × 64.8 µm^2^.

**Video S3. Close-up view of example particle merging and splitting events for Halo-FLR2**. This example is from the movie shown in Video S1 and corresponds to the example shown in Fig. 1D. Shown are the raw movie (top) and the particle tracks overlaid on the movie (bottom). In this example, a particle splits into two particles at 11.6 s, which move separately for some tie, and then re-merge with each other at 12.6 s. The merging/splitting events are indicated by yellow/green diamonds at the time points just before and just after each event. The detections at all other time points are shown as red circles, with tracks shown in light pink. Image size is 4.1 × 4.1 µm^2^.

**Video S4. Close-up view of example particle merging and splitting events for ECTM-Halo**. This example is from the movie shown in Video S2. Shown are the raw movie (top) and the particle tracks overlaid on the movie (bottom). In this example, two particles merge at 1.6 s, stay together for some time, and then split apart at 2.9 s. The merging/splitting events are indicated by yellow/green diamonds at the time points just before and just after each event. The detections at all other time points are shown as red circles, with tracks shown in light pink. Image size is 4.1 × 4.1 µm^2^.

**Video S5. Synthetic single-molecule movie**. Representative 10 Hz/30 s synthetic movie, using the diffusion coefficient, density and labeled fraction parameters of FLR2. This simulation shows an example of no interactions. Shown are the raw movie (top) and the particle tracks overlaid on the movie (bottom). Tracks are color-coded randomly with 7 different colors for ease of visualization. Image size is 27.5 × 27.5 µm^2^.

## Acknowledgments

We thank Dr. Tieqiao Zhang for microscopy support, and Joseph Chi (Danuser lab, UT Southwestern) for help with making the ECTM-Halo construct. This work was supported by funding from the National Institutes of Health/National Institute of General Medical Sciences (R35 GM119619), the Welch Foundation (I-1901-20190330), and the University of Texas Southwestern Medical Center Endowed Scholars Program to K. Jaqaman. J. Guerrero was a trainee of the National Institutes of Health Molecular Biophysics Training Grant (5T32GM131963; PI: Dr. Yuh Min Chook).

## Supplementary Notes

### Supplementary Note S1: Step photobleaching analysis

In the single-molecule regime, fluorophore photobleaching is captured as discrete jumps or steps in particle intensity traces over time, ultimately resulting in particle disappearance. Thus, the photobleaching step sizes reflect the intensity distribution of an individual fluorophore. In addition, for intensity traces that disappear before the end of the SMI movie, the number of photobleaching steps within the trace + the disappearance step reflects the number of fluorophores in the particle. In other words, a trace with n photobleaching steps within it represents n+1 fluorophores, bleaching one at a time. Therefore, the average intensity of each trace segment between consecutive photobleaching steps, and between the last photobleaching step and disappearance, represents the intensity of a specific number of fluorophores, depending on the number of steps until disappearance.

We performed photobleaching analysis as follows: Each intensity trace (from a particle track) was fitted with a multistep function using least squares fitting. The number of steps within the trace was determined by fitting first with 0 steps, then with 1 step, then with 2 steps, etc. and comparing the residuals from employing n+1 steps to those from employing n steps using an F-test. If employing n+1 steps was found to significantly improve the fit when compared to n steps (using an α-value of 0.05), then n+1 steps were accepted and n+2 steps were attempted, and so on and so forth. This nested model selection strategy yielded the minimum number of steps necessary to fit each intensity trace and avoid overfitting [REF]. Once the number of steps for a particular trace was determined, the model fit yielded the locations of the different steps and their sizes, as well as the average intensity of the intensity trace in between the detected photobleaching steps, and between the last photobleaching step and disappearance. See Fig. S2A for examples of multistep fitting to intensity traces.

We applied step photobleaching analysis to low labeling density SMI data. The reasons were twofold: (1) Low labeling reduced the chances of particle merging and splitting, thus simplifying the acquired particle tracks and intensity traces. (2) Having much less particles in the field of view than the regular SMI movies, low labeling increased tracking accuracy, thus increasing our confidence in the particle tracks and intensity time traces used for photobleaching analysis.

Step photobleaching analysis of these data yielded the distribution of photobleaching step sizes, reflecting the distribution of individual JF549 fluorophore intensities, in our imaging set up (Fig. S2B). It also yielded the average intensities of the trace segments in between photobleaching steps, or between the last photobleaching step and disappearance, grouped by their number of steps until disappearance (Fig. S2C, D). Note that the average trace segment intensity just before disappearance (blue histograms in Fig. 2C, D), which represented one fluorophore, spanned the same intensity range as the histogram of photobleaching step sizes (which also represented one fluorophore). This analysis revealed that the intensity of individual fluorophores in our imaging set up spanned a wide range, resulting in large overlap in intensity between particles containing different numbers of fluorophores.

Because of this large overlap and difficulty in mapping particle intensity back to number of fluorophores, i.e. apparent oligomeric data of a detected particle, intensity was not used in the process of determining the oligomeric state. Note, however, that although the label was extracellular in the case Halo-FLR2 and intracellular in the case of ECTM-Halo, their single fluorophore intensity properties were similar.

### Supplementary Note S2: Analytical mapping between full populations and labeled subsets of receptors and receptor complexes

#### Full population of receptors and receptor complexes

Let there be a receptor species that can get organized into complexes, which can be monomers (1 receptor alone), dimers (2 receptors together), trimers (3 receptors together), etc. Let *n* be the oligomeric state of a complex and *N* be the maximum oligomeric state achievable. Let 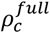 be the total density of receptor complexes regardless of their oligomeric state, and 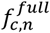 be the fraction of complexes with oligomeric state *n*. Then, the density of receptor complexes with oligomeric state 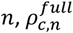, is given by:

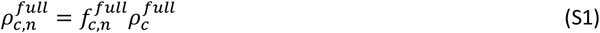

Note that 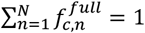 and

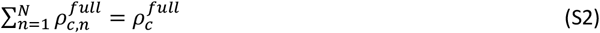

Note also that the density of receptors 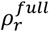 and the density/fraction of complexes are related to each other as follows:

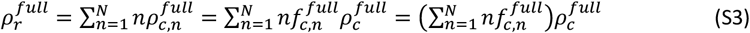

#### Labeled subset of receptors and receptor complexes

When only a fraction of the receptors is labeled, a receptor complex with none of its receptors labeled will not be observed. In addition, among the observed complexes, the apparent oligomeric state of a complex can shift toward smaller values, depending on how many receptors in that complex happen to be labeled. True monomers, when observed, can only appear as monomers in the labeled subset. True dimers, however, can appear as monomers (if only 1 of the 2 receptors is labeled) or dimers (if both receptors are labeled). True trimers can appear as monomers, dimers, or trimers. Etc. Thus, the distribution of oligomeric states among the observed complexes will shift in the labeled subset. From the perspective of the labeled (i.e. observed) subset of complexes, apparent monomers can originate from true monomers, true dimers, true trimers, etc. Apparent dimers can originate from true dimers, true trimers, etc. And so on and so forth.

Define 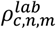 as the density of labeled complexes with apparent oligomeric state *n*, originating from full population complexes with true oligomeric state *m*. The relationship between 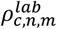 and 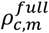 is derived from the binomial distribution 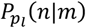, with *p*_*l*_ = the probability of success = the probability of a receptor getting labeled, *m* = number of Bernoulli trials and *n* = number of successes (i.e., receptors labeled):

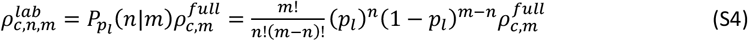

With this, the total density of labeled complexes with apparent oligomeric state *n* regardless of the true oligomeric state of the complexes that they originate from, 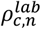, is given by:

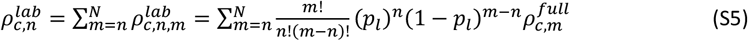

Moving forward, it is useful to substitute 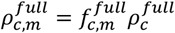 (from Eq. S1) in Eq. S4 and rewrite Eq. S4 as:

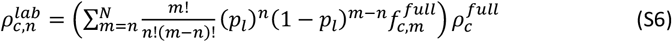

The total density of labeled receptor complexes regardless of their apparent oligomeric state, 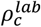, is then given by:

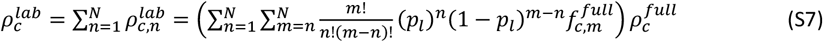

Eqs. S4-S7 allow us to write the fraction of labeled complexes with apparent oligomeric state *n* and the fraction of receptor complexes that gets labeled, i.e. observed, regardless of their apparent oligomeric state, as functions of the labeled fraction of receptors *p*_*l*_ and the fraction of complexes of different oligomeric states in the full population 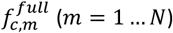. Specifically, dividing Eq. S6 by Eq. S7 yields the fraction of labeled complexes with apparent oligomeric state 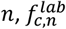:

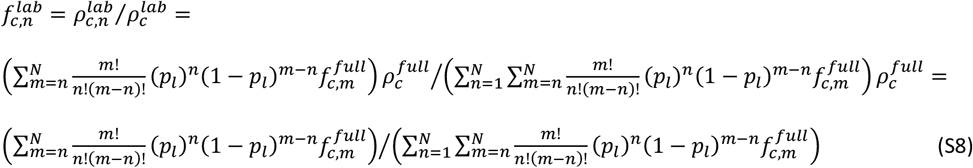

Similarly, dividing Eq. S7 by the density of receptor complexes in the full population, 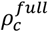, yields the fraction *q*_*l*_ of receptor complexes that gets labeled, i.e. observed. As a reminder, a complex is observed as long as at least one of its subunits gets labeled.

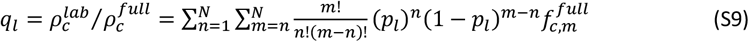

## Supplementary Figures

**Figure S1.**
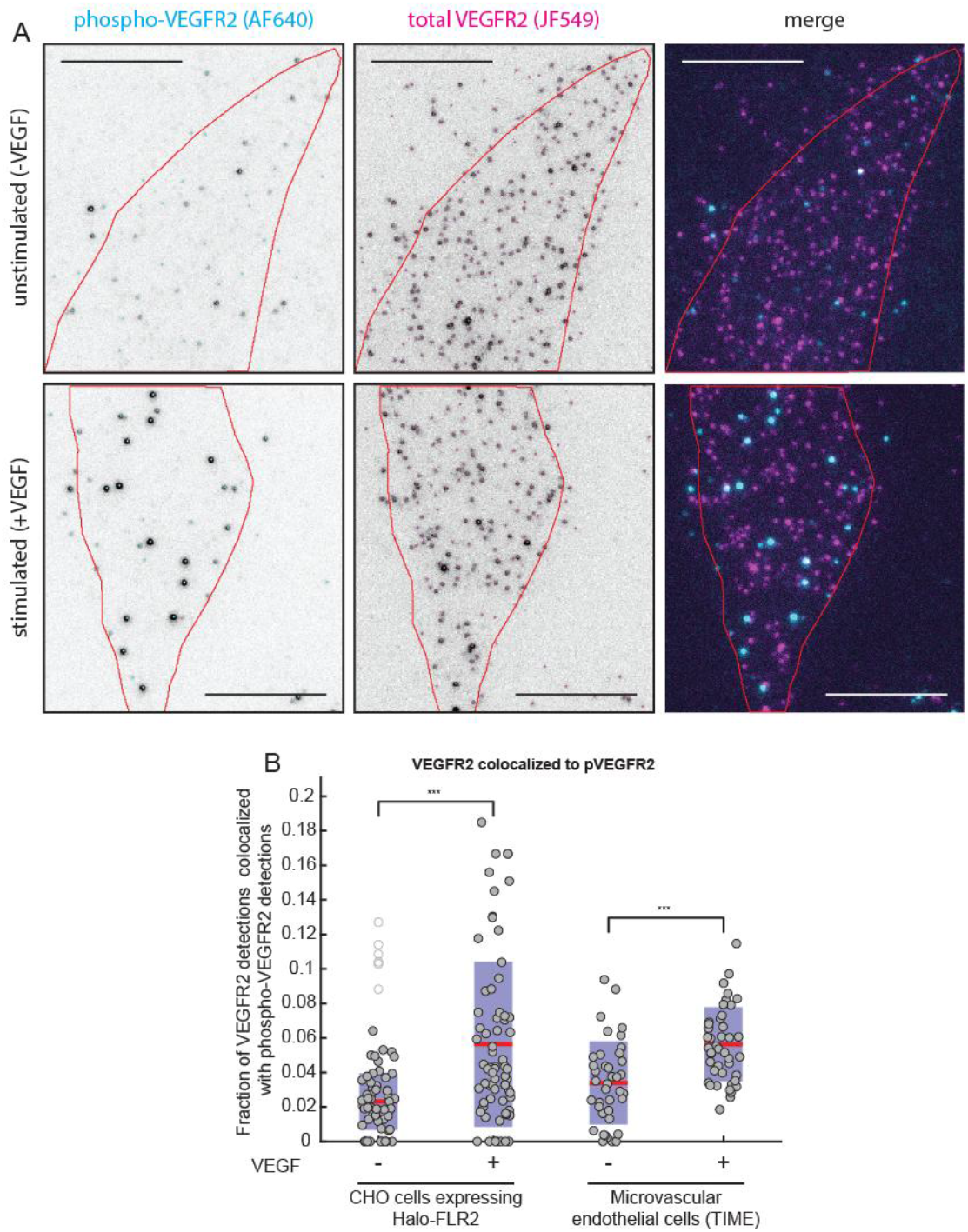
Validation of Halo-FLR2 construct. **(A)** Representative 2-color fixed-cell image of Halo-FLR2 in CHO cells labeled with Halo-ligand, together with an anti-phospho-VEGFR2 antibody, in the absence (top) or presence of 2 nM VEGF for 5 min. Cell outline is shown in red. The individual channel images are inverted for visual clarity. Scale bar, 10 µm. **(B)** Fraction of VEGFR2 detections (Halo-FLR2 in CHO cells or endogenous VEGFR2 in TIME cells) colocalized with the anti-phospho-VEGFR2 antibody, as a measure of VEFGR2 phosphorylation in the absence or presence of VEGF. Circles indicate individual cell measurements, with open circles indicating outliers. Red lines and shaded bars show mean and standard deviation, respectively, over group of cells. ***, p < 0.001, where -VEGF and +VEGF measurements were compared using a 2-sample t-test. The rise in Halo-FLR2 phosphorylation in CHO cells in response to VEGF is similar to that of endogenous VEGFR2 in TIME cells, indicating that Halo-FLR2 is able to bind VEGF and to get subsequently activated. Dataset consisted of 60, 69, 39 and 44 cells from 9, 10, 4 and 4 independent repeats for CHO -VEGF, CHO +VEGF, TIME -VEGF and TIME +VEGF, respectively.

**Figure S2.**
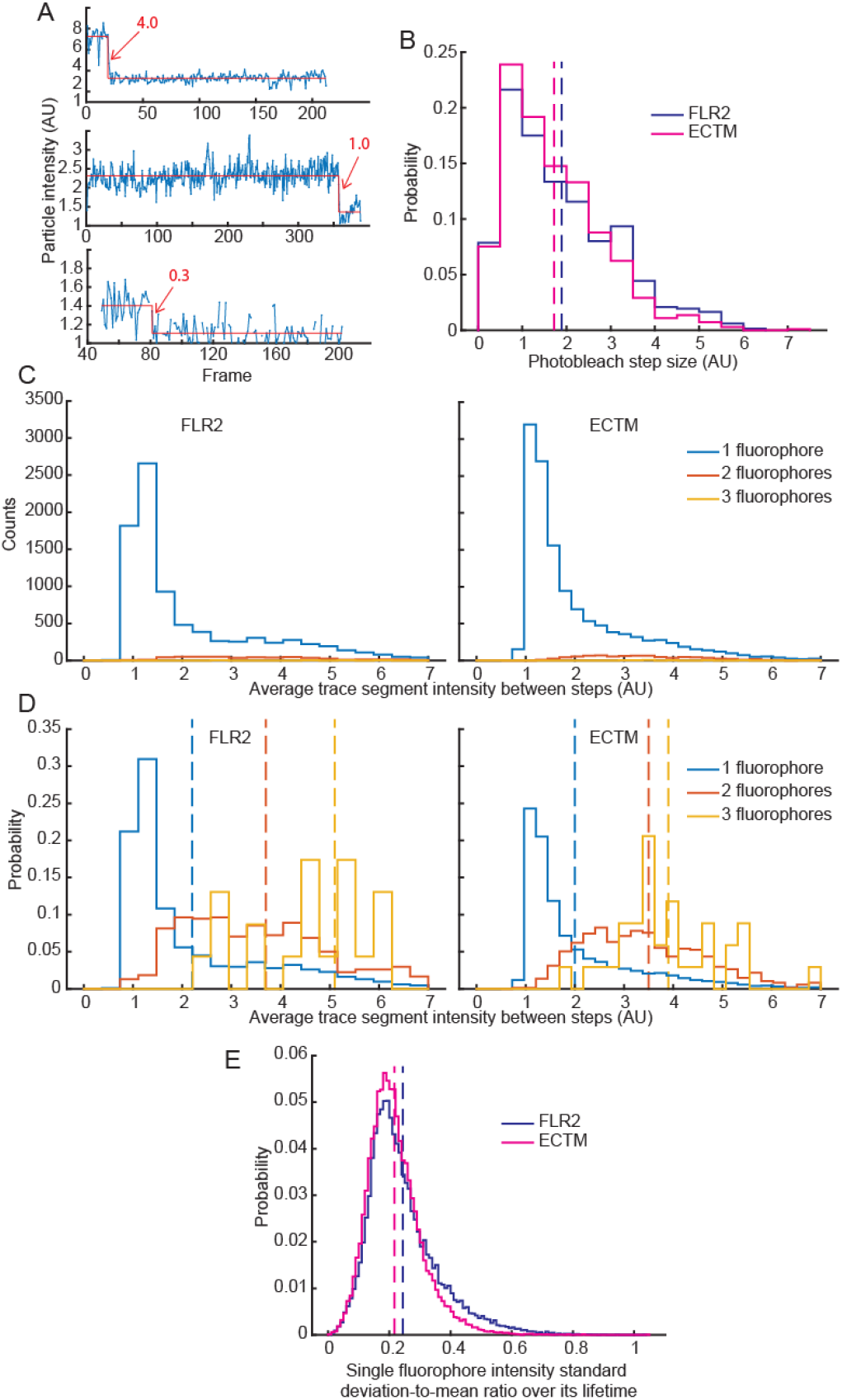
Analysis of fluorophore intensity properties in SMI data. **(A)** Example photobleaching steps for three particles. Each of these particles photobleaches once in its lifetime, at frames 20 (top), 358 (middle) and 81 (bottom), and then disappears sometime later (due to another photobleaching step, but which is not explicitly detected). Values in red indicate step size. **(B)** Distribution of photobleaching step sizes for FLR2 (blue) and ECTM (magenta), from the photobleaching steps that do not lead to particle disappearance (because particle disappearance could be due to photobleaching or tracking errors). Dashed vertical lines show the mean value. **(C, D)** Distribution of average particle intensity between photobleaching steps, shown as counts **(C)** and probability **(D)**, for FLR2 (left) and ECTM (right). Blue, red and yellow represent different categories of particles, specifically those that are, respectively, 1, 2 and 3 photobleaching steps away from disappearance. The histograms as counts in C highlight the dominance of the 1-step-to-photobleaching category (i.e., apparent monomers). The histograms as probabilities in D highlight the overlap in particle intensities between the different categories. Vertical dashed lines show mean values of intensity distributions for 1 fluorophore (2.2 for FLRs and 2.0 for ECTM), 2 fluorophores (3.7 for FLR2 and 3.5 for ECTM) and 3 fluorophores (5.1 for FLR2 and 3.9 for ECTM). The mean intensity over all particles of any particular category correlates with the number of fluorescing fluorophores in that particle, as expected. However, the intensity of any individual particle in a category can be far from the mean, leading to the high overlap between particles of different categories. **(E)** Single fluorophore intensity standard deviation-to-mean ratio, calculated per fluorophore over its lifetime. Only particles with mean intensity compatible with a single fluorophore were used for this analysis (see Materials and Methods). In B-E, distributions are over all detected particles or events (as relevant) over all cells employed in analysis. For A-D, dataset consisted of 39 (FLR2) and 48 (ECTM) cells, combined from 5 (FLR2) and 5 (ECTM) independent repeats of SMI with low labeling density (JF549 at 0.5 nM for FLR2 and 1 nM for ECTM). For E, dataset was same as in Fig. 1.

**Figure S3.**
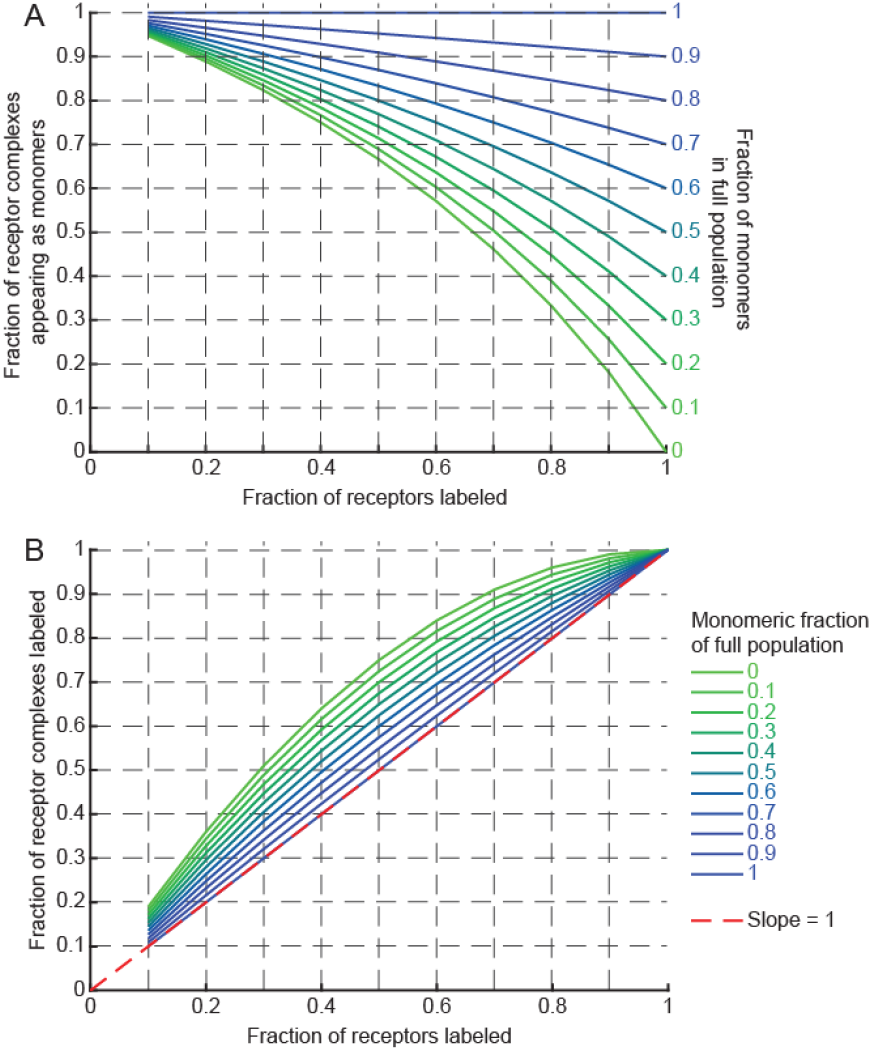
Analytical mapping between full population and labeled subset in a system that contains monomers and dimers. **(A)** Fraction of apparent monomers among the labeled subset of complexes as a function of the fraction of receptors that are labeled, for systems with indicated true fractions of monomers among the full population of complexes. As the fraction of labeled receptors decreases, the fraction of apparent monomers among the labeled complexes increases. **(B)** Fraction of complexes that get labeled as a function of the fraction of receptors that are labeled, for systems with indicated true fractions of monomers among the full population of complexes. As the fraction of monomers in the full system decreases (i.e., the fraction of dimers increases), the fraction of complexes that get labeled deviates further from the fraction of receptors that are labeled. The fraction of complexes that get labeled ≥ the fraction of receptors that are labeled under all conditions.

**Figure S4.**
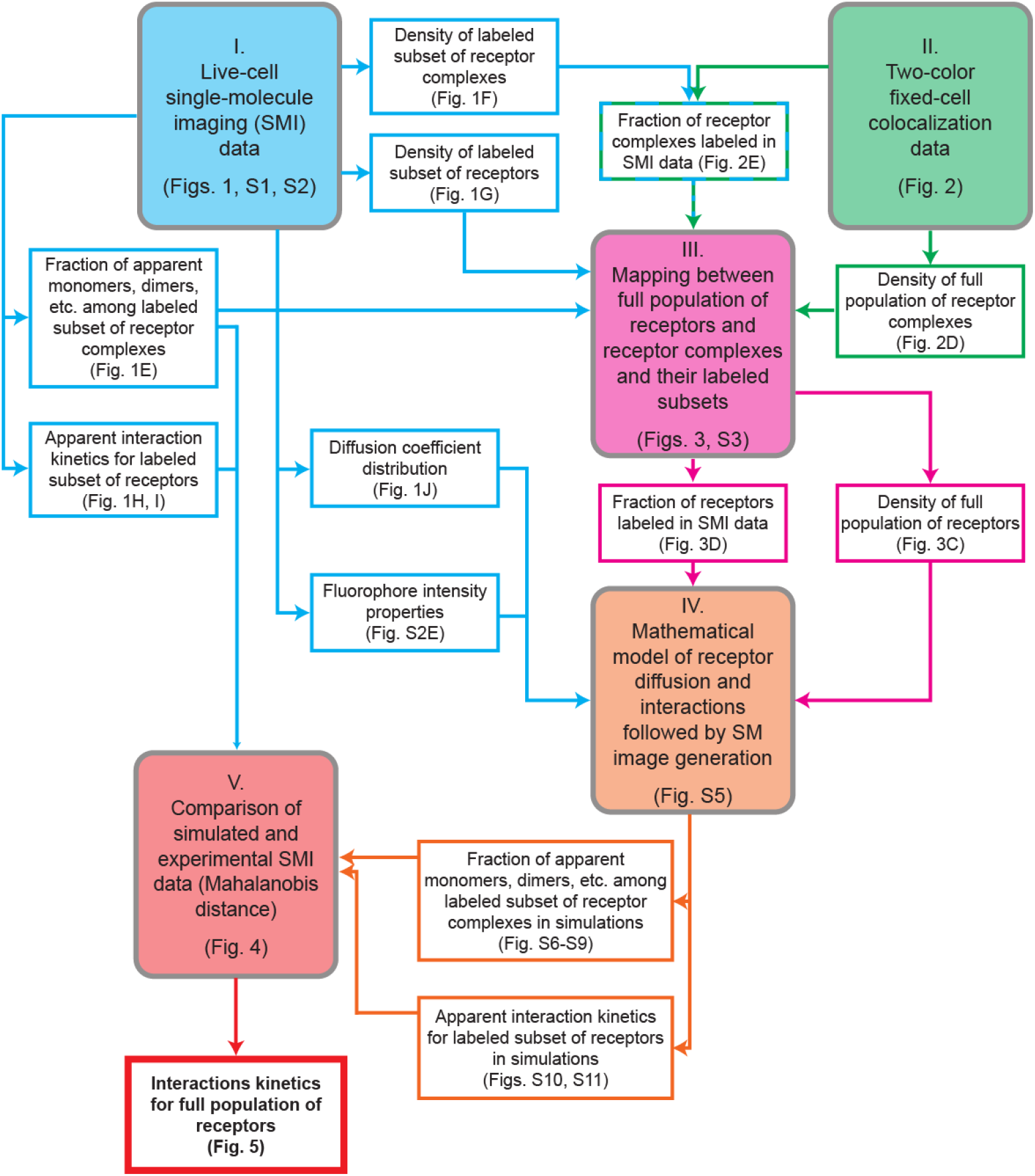
Flowchart of experiments, analyses and data flow to determine the kinetics of VEGFR2 homotypic interactions on the cell surface. The workflow consists of five blocks, each represented by a different color. Blocks I and II represent the major experimental legs of the workflow and data derived from them. Blocks III and IV represent the major analytical and computational legs and the data derived from them. Block V represents the comparison of experimental and simulated data to derive the interaction kinetics parameters of interest. The flowchart indicates the figures/figure panels belonging to each block and its data.

**Figure S5.**
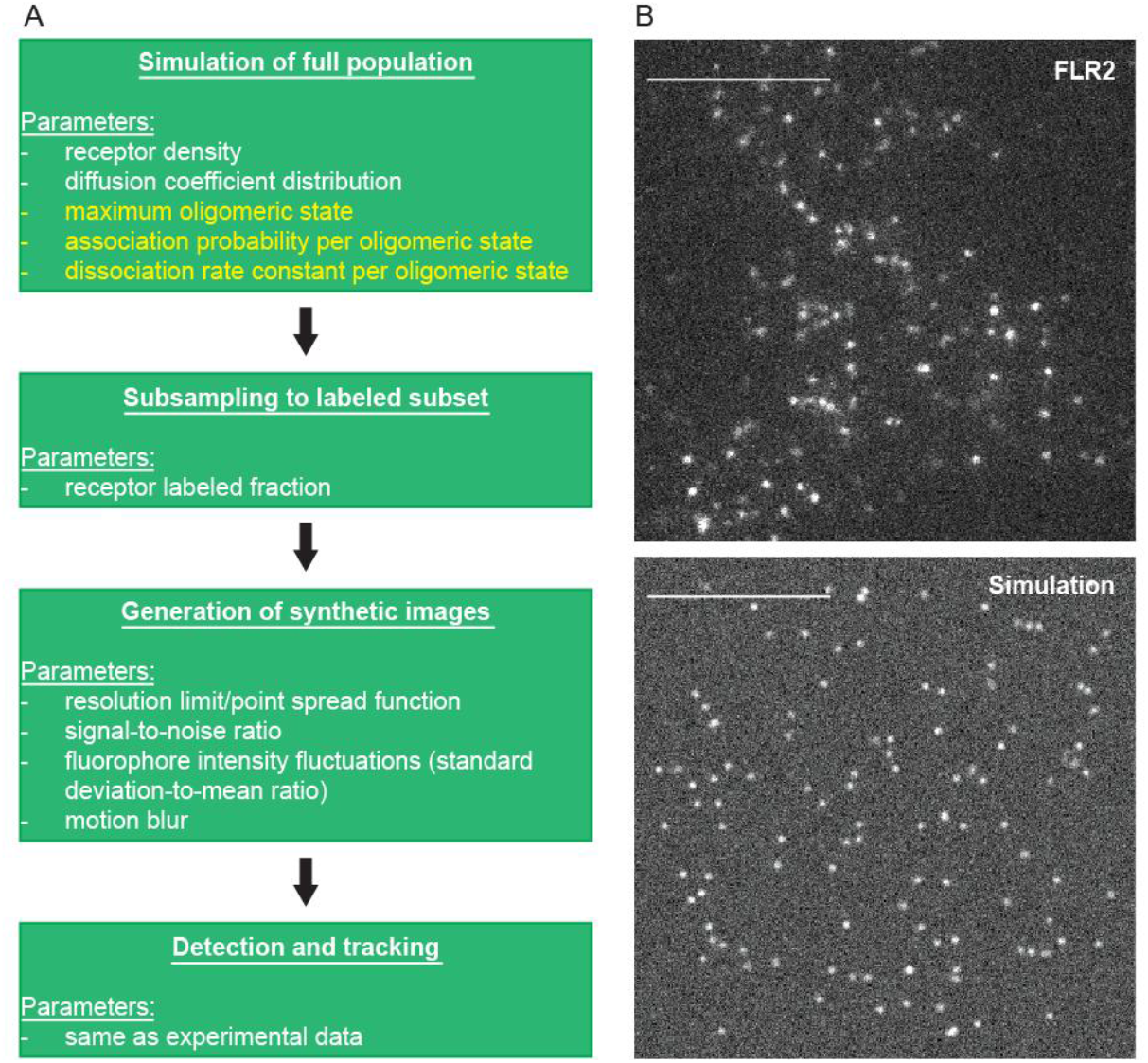
Pipeline of generating synthetic single-molecule data of VEGFR2 diffusion and interactions to compare to experimental SMI data. **(A)** The four steps of the pipeline, with the parameters involved in each step. Parameters in white are derived directly from experimental data. Parameters in yellow are determined by comparing simulated data to experimental data. **(B)** Representative real (top) and simulated (bottom) images, for visual comparison. Scale bar, 10 μm.

**Figure S6.**
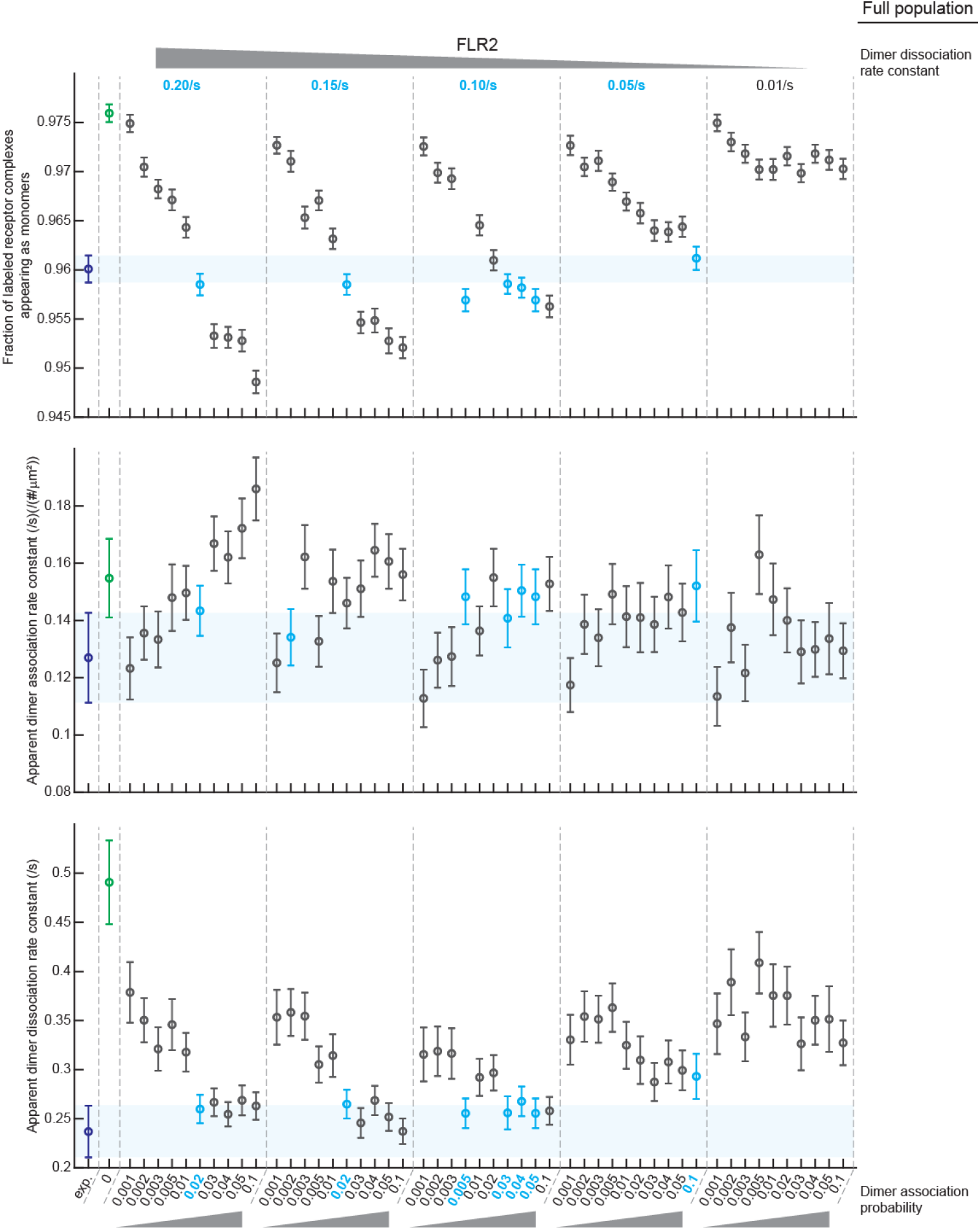
Statistics describing experimental and simulated FLR2 interactions, Part I. Fraction of labeled receptor complexes appearing as monomers (top), apparent dimer association rate constant (middle), and apparent dimer dissociation rate constant (bottom). For all, the left-most data point (blue) is for experimental data, the second data point (green) is for simulations of no interactions [does not match experimental data], while the remaining data points (black [not matching experimental data] and cyan [matching experimental data]) belong to a series of dimerization simulations with the indicated full population dimer association probability and dissociation rate constant. The simulation results are grouped by dissociation rate constant, within each of which the association probability is varied as indicated. The different groups are delineated by dashed vertical lines. Simulations are considered to match the experimental data if their Mahalanobis distance to the simulated data < 700 (see Fig. 4) and are highlighted by coloring their circles and parameter values in cyan. Circles show fractions from analyzing all imaged cells or simulations together. Error bars show the standard deviation of the intermediate statistics, estimated using Jackknife resampling. Shaded rectangle in the background of each plot shows the mean ± 1 standard deviation for the experimental data, for visual comparison to the simulated data. Experimental intermediate statistics values are from Fig. 1. Simulated intermediate statistics are from 200 simulations for each parameter combination.

**Figure S7.**
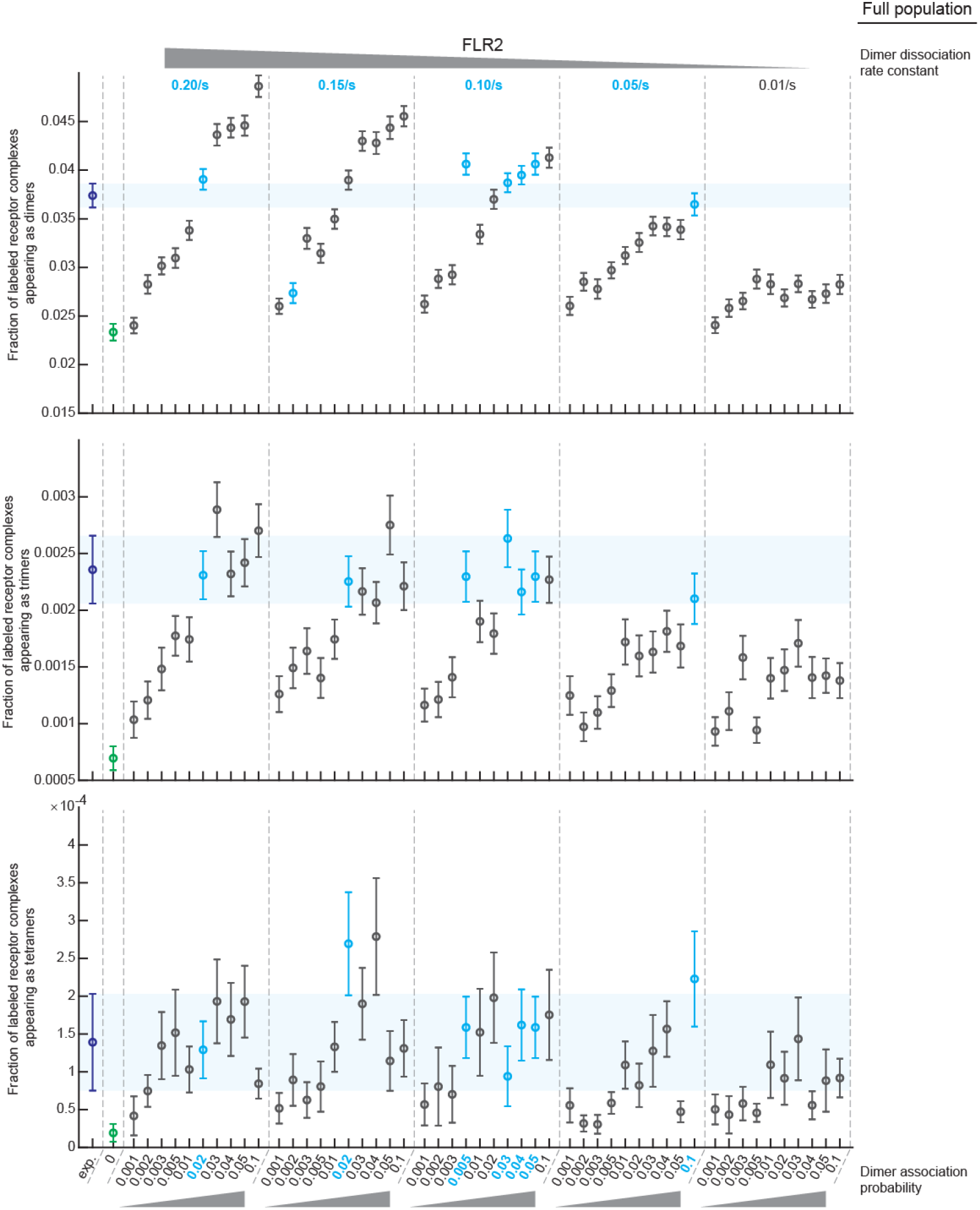
Statistics describing experimental and simulated FLR2 interactions, Part II. Same as Fig. S6, but for the fraction of labeled receptors appearing as dimers (top), trimers (middle), and tetramers (bottom). All details as in Fig. S6.

**Figure S8.**
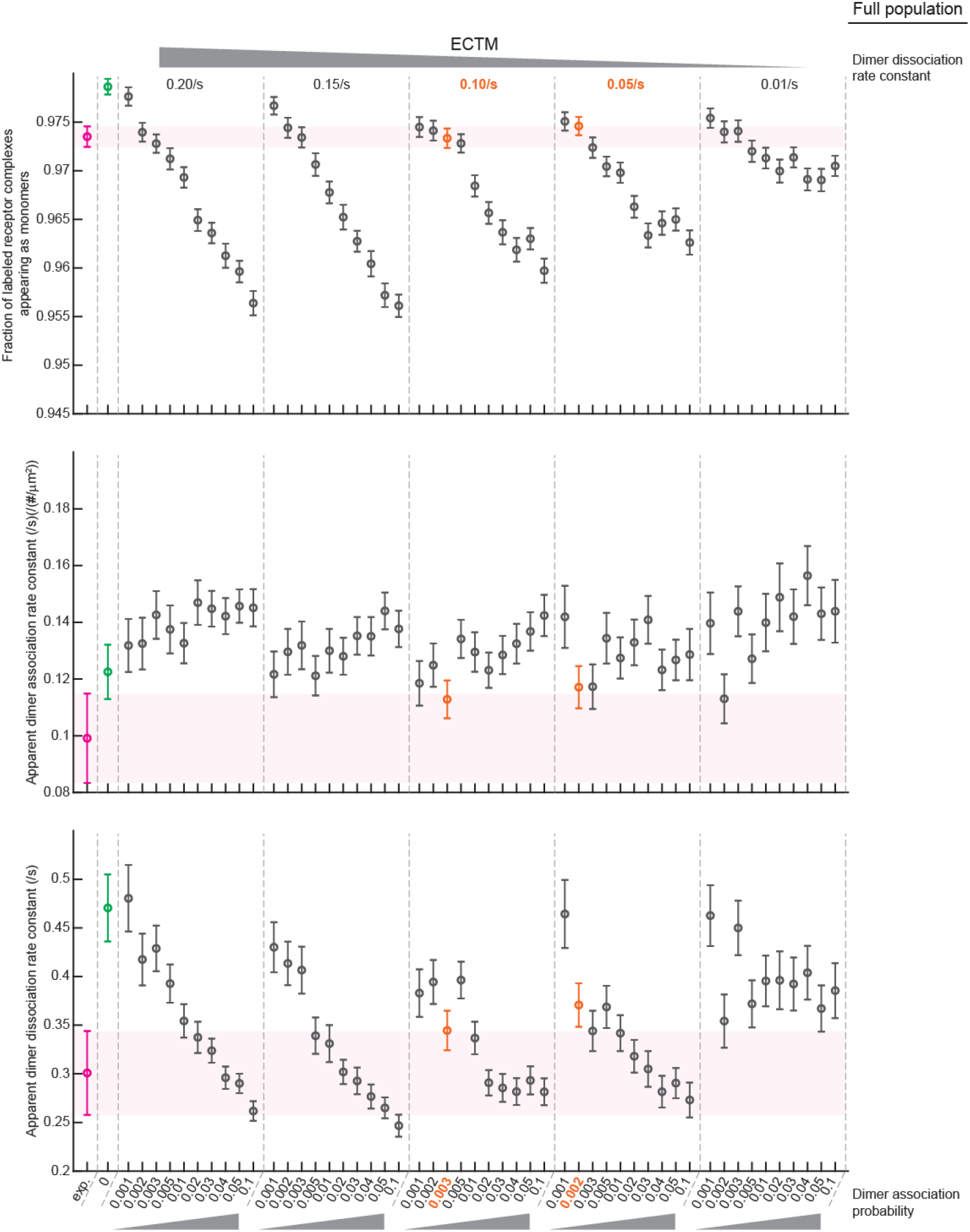
Statistics describing experimental and simulated ECTM interactions, Part I. Fraction of labeled receptor complexes appearing as monomers (top), apparent dimer association rate constant (middle), and apparent dimer dissociation rate constant (bottom). All details as in Fig. S6, except that experimental data points are in magenta and simulated data points matching experimental data are in orange.

**Figure S9.**
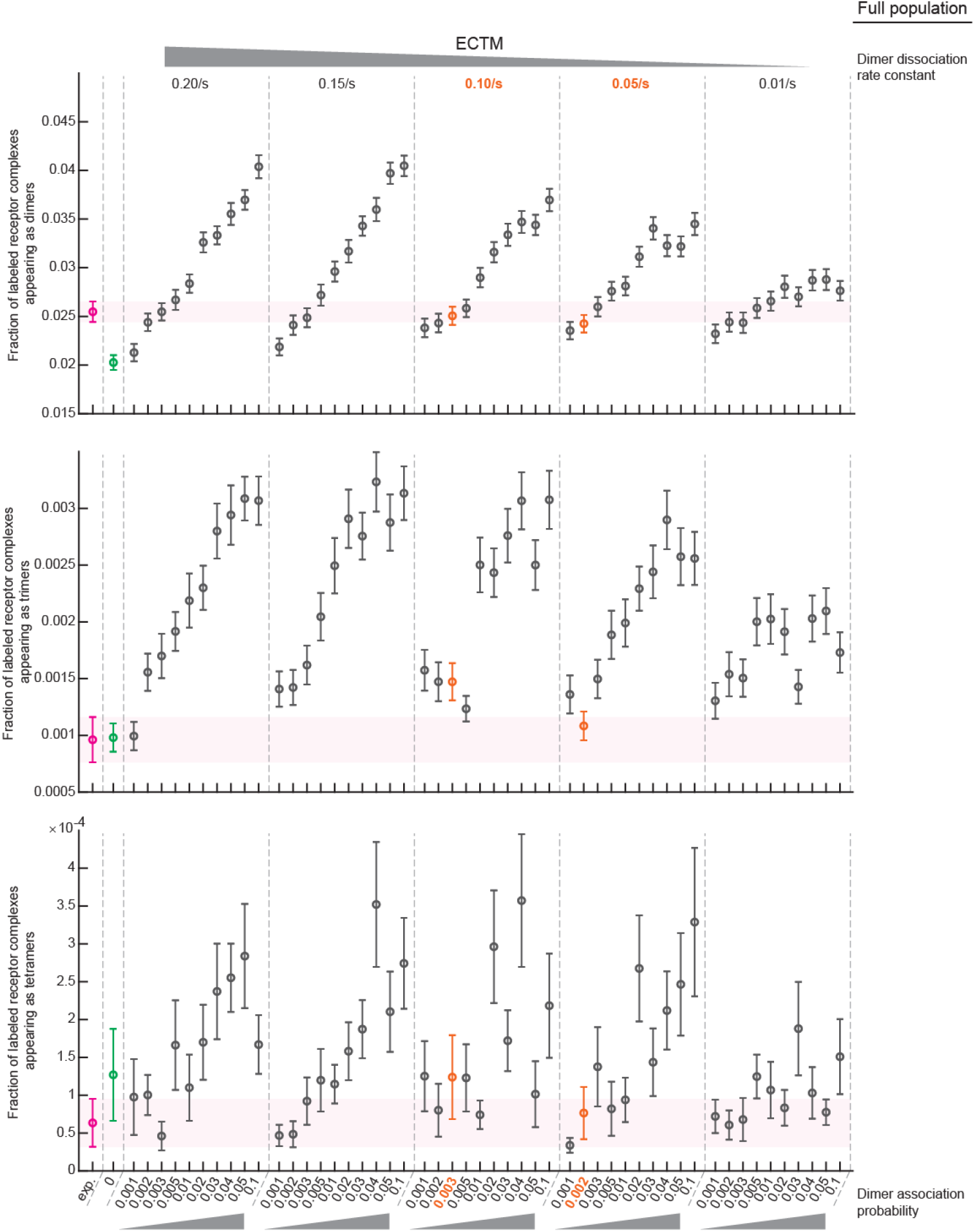
Statistics describing experimental and simulated ECTM interactions, Part II. Same as Fig. S8, but for the fraction of labeled receptors appearing as dimers (top), trimers (middle), and tetramers (bottom). All details as in Fig. S8.

